# Transcriptional profiling reveals potential involvement of microvillous TRPM5-expressing cells in viral infection of the olfactory epithelium

**DOI:** 10.1101/2020.05.14.096016

**Authors:** B. Dnate’ Baxter, Eric D. Larson, Laetitia Merle, Paul Feinstein, Arianna Gentile Polese, Andrew N. Bubak, Christy S. Niemeyer, James Hassell, Doug Shepherd, Vijay R. Ramakrishnan, Maria A. Nagel, Diego Restrepo

**Author notes:** Co-first authors (B. Dnate’ Baxter, Eric D. Larson, Laetitia Merle). Corresponding author: Diego Restrepo.

## Abstract

**Background:** Understanding viral infection of the olfactory epithelium is essential because the olfactory nerve is an important route of entry for viruses to the central nervous system. Specialized chemosensory epithelial cells that express the transient receptor potential cation channel subfamily M member 5 (TRPM5) are found throughout the airways and intestinal epithelium and are involved in responses to viral infection.

**Results:** Herein we performed deep transcriptional profiling of olfactory epithelial cells sorted by flow cytometry based on the expression of mCherry as a marker for olfactory sensory neurons and for eGFP in OMP-H2B::mCherry/TRPM5-eGFP transgenic mice (*Mus musculus*). We find profuse expression of transcripts involved in inflammation, immunity and viral infection in TRPM5-expressing microvillous cells.

**Conclusion:** Our study provides new insights into a potential role for TRPM5-expressing microvillous cells in viral infection of the olfactory epithelium. We find that, as found for solitary chemosensory cells (SCCs) and brush cells in the airway epithelium, and for tuft cells in the intestine, the transcriptome of TRPM5-expressing microvillous cells indicates that they are likely involved in the inflammatory response elicited by viral infection of the olfactory epithelium.

## Background

Chemosensory cells found in the airway (SCCs/brush cells) and intestinal epithelium (tuft cells) express the transient receptor potential cation channel subfamily M member 5 (TRPM5) and other elements of the taste transduction pathway and have been implicated in immune and inflammatory responses to bacterial, viral and parasitic infection (Luo et al., 2019; Maina et al., 2018; O’Leary et al., 2019; Perniss et al., 2020; Rane et al., 2019; Saunders et al., 2014; Tizzano et al., 2010). In the olfactory epithelium TRPM5 and other proteins involved in taste transduction are also expressed in SCC-like microvillous cells (MVCs)(Genovese and Tizzano, 2018; Lin et al., 2008), which have been proposed to be involved in a protective response to high concentrations of odorants (Fu et al., 2018; Lemons et al., 2017). However, whether MVCs play a role in viral infection or viral infection defense of the olfactory epithelium is unknown.

Herein, we performed transcriptional profiling of MVCs and a subset of olfactory sensory neurons (OSNs) expressing eGFP under control of a fragment of the TRPM5 promoter (OSN_eGFP+ cells)(Lin et al., 2007; Lopez et al., 2014). In order to profile these low abundance cells we used a modified version of Probe-Seq, which allows deep transcriptional profiling of specific cell types identified by fluorescent markers as the defining feature (Amamoto et al., 2019). We crossed a mouse expressing mCherry in the nuclei of OSNs under control of the OMP promoter (OMP-H2B::mCherry mice) with TRPM5-eGFP transgenic mice (Clapp et al., 2006) (OMP-H2B::mCherry/TRPM5-eGFP mice). We isolated cells from the olfactory epithelium and used fluorescence-activated cell sorting (FACS) to sort MVC_eGFP cells (mCherry negative and eGFP positive) and cells labeled by OMP-driven mCherry that did or did not express eGFP (OSN_eGFP+ and OSN_eGFP-cells) followed by transcriptional profiling by RNA sequencing (RNAseq).

## Results

### Fluorescence-activated cell sorting of cells isolated from the main olfactory epithelium

The olfactory epithelium of OMP-H2B::mCherry/TRPM5-eGFP mice expressed nuclear mCherry driven by the OMP promoter in the intermediate layer of the olfactory epithelium (Figure 1a), as expected for the location of nuclei of mature OSNs (Farbman and Margolis, 1980). eGFP expression driven by the TRPM5 promoter was found in MVCs, with cell bodies located mostly in the apical layer of the epithelium (asterisks), and at lower expression levels in a subset of OSNs double-labeled with mCherry (Figure 1a), consistent with earlier publications (Lin et al., 2008; Lin et al., 2007).

**Figure 1.**
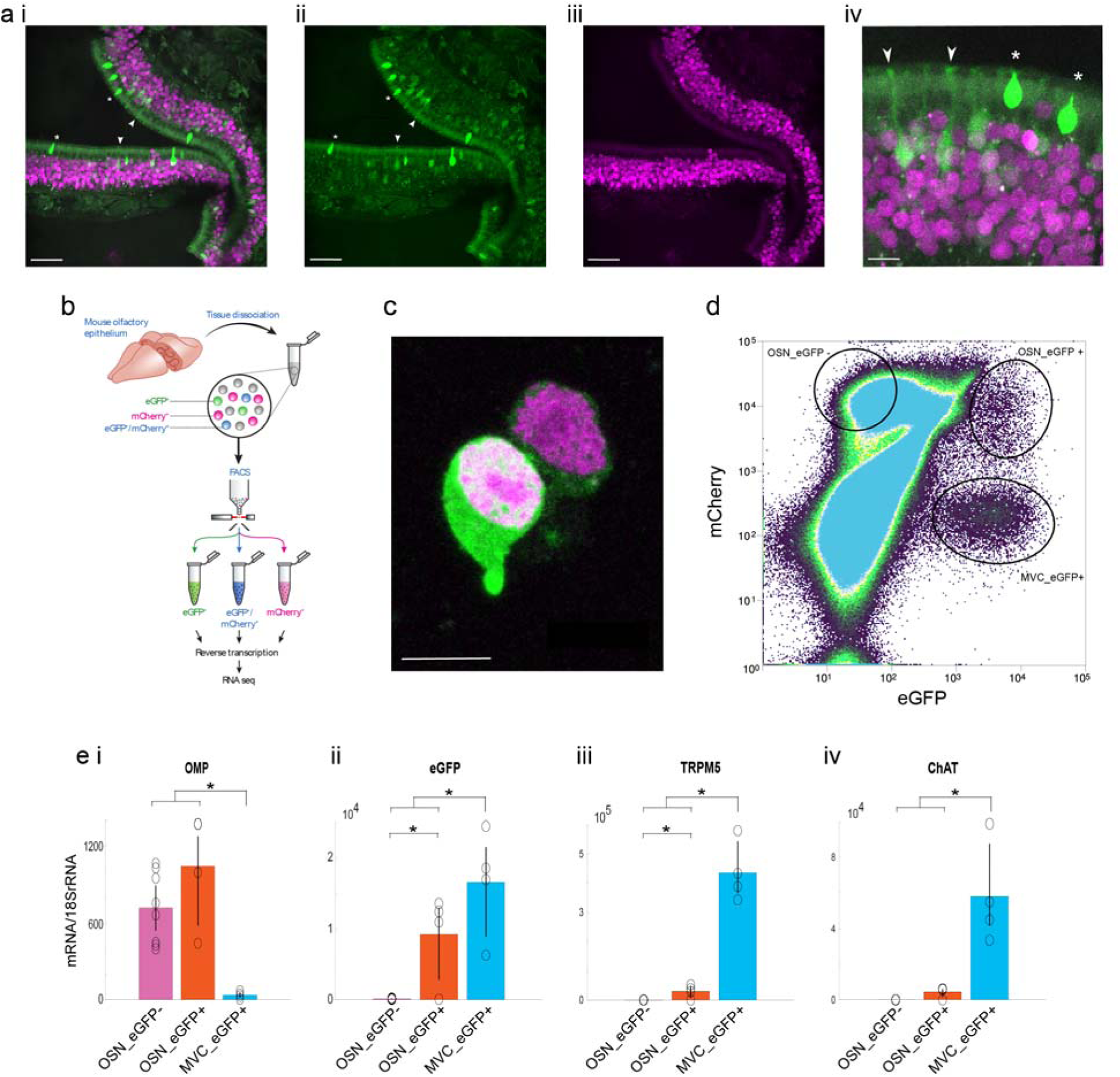
Fluorescence activated sorting (FACS) of cells isolated from the olfactory epithelium. **a.** TRPM5 promoter driven expression of eGFP and OMP promoter driven expression of mCherry in the olfactory epithelium. Expression of eGFP is found both in MVCs that do not express mCherry (asterisk) and in OSNs double labeled with eGFP and mCherry (arrow). **i.** Composite, **ii.** eGFP, **iii.** mCherry, **iv**. Composite magnification. Magenta: mCherry, green: eGFP. Scale bar: i-iii, 50 μm, iv, 10 μm. **b.** Schematic of RNA-seq process from tissue to RNA extraction. Mouse OE was dissociated into single cells and sorted via FACS. RNA was extracted from each of the resulting cell populations. **c.** Two isolated OSNs differing in eGFP expression. Magenta: mCherry, green: eGFP. Scale bar: 10 μm. **d.** Distribution of mCherry and eGFP fluorescence intensity for FACS-sorted cells. Three cell populations were isolated for RNAseq: Cells with low OMP promoter-driven mCherry expression and high TRPM5 promoter-driven eGFP expression (MVC_eGFP cells), cells with high OMP promoter-driven mCherry and low eGFP expression (OSN_eGFP- cells) and cells with eGFP expression of the same magnitude as MVC_eGFP cells and high OMP promoter- driven mCherry expression (OSN_eGFP+ cells). The number of cells collected for this FACS run were: OSN_eGFP-s 1,500,000, OSN_eGFP+s 5336 and MVC_eGFP cells 37,178. **e.** qPCR levels (normalized to levels 18s RNA) for expression of transcripts encoding for OMP (**i**), TRPM5 (**ii**), eGFP (**iii**) and ChAT (**iv**). The asterisks denote significant differences tested with either t-test or ranksum with p-values below the significance p-value corrected for multiple comparisons using the false discovery rate (pFDR)(Curran-Everett, 2000). pFDR is 0.033 for OMP, 0.05 for TRPM5, 0.05 for eGFP and 0.03 for ChAT, n=8 for OMP OSN_eGFP-s, 4 for OMP OSN_eGFP+s and 4 for MVC_eGFP cells.

We proceeded to isolate cells from the main olfactory epithelium of OMP-H2B::mCherry/TRPM5-eGFP mice (see Figure 1b, Methods and Figure 1 - figure supplement 1). Figure 1c shows two isolated OSNs with differential expression of eGFP. Using flow cytometry we found that fluorescence intensity of individual cells for mCherry and eGFP spanned several orders of magnitude (Figure 1d). We proceeded to sort three groups of cells under light scattering settings to exclude doublets: high mCherry-expressing cells with low and high eGFP fluorescence (presumably mature OSNs, these cells are termed OSN_eGFP- and OSN_eGFP+ cells respectively) and cells with low mCherry and high eGFP expression (MVC_eGFP, presumably MVCs). Reverse transcription quantitative PCR (RT-qPCR) showed that, as expected the OSN_eGFP- and OSN_eGFP+ cells have higher levels of OMP transcript than MVC_eGFP cells (Figure 1e,i), and OSN_eGFP+ cells and MVC_eGFP cells have higher levels of eGFP transcript compared to OSN_eGFP- cells (Figure 1e,ii). Furthermore, compared to OSN_eGFP- cells both the MVC_eGFP cells and OSN_eGFP+ cells expressed higher levels of TRPM5 transcript (Figure 1e,iii) and choline acetyl transferase (ChAT)(Figure 1e,iv), a protein involved in acetylcholine neurotransmission that is expressed in MVCs (Ogura et al., 2011). The asterisks in Figure 1e denote significant differences tested with either t-test or ranksum with p-values below the p-value of significance corrected for multiple comparisons using the false discovery rate (pFDR)(Curran-Everett, 2000) (pFDR is 0.033 for OMP, 0.05 for TRPM5, 0.05 for EGFP and 0.03 for ChAT, n=8 for OMP OSN_eGFP-, 4 for OMP OSN_eGFP+ and 4 for MVC_eGFP cells).

### RNAseq indicates that MVC_eGFP and OSN_eGFP- are distinct groups of chemosensory cells in the mouse olfactory epithelium

Differential gene expression analysis of the RNAseq data was used to compare MVC_eGFP and OSN_eGFP- sorted by FACS. Expression of 4386 genes was significantly higher in MVC_eGFP cells compared to OSN_eGFP- cells, and expression of 5630 genes was lower in MVC_eGFP cells (Figure 2a shows the most significantly upregulated or downregulated genes and Figure 2 – figure supplement 1 shows the entire list). A total of 1073 Olfr genes were included in the analysis (including pseudogenes). While 9 were expressed at higher levels in the MVC_eGFP population their expression levels was very low (<100 counts) and the difference was not statistically significant. On the contrary, transcripts for 550 olfactory receptors were significantly higher in OSN_eGFP- cells (Figure 2b and 2c). *Trpm5* and *eGFP* were among the top 10 genes whose transcription was higher in MVC_eGFP cells compared to OSN_eGFP- cells with 1471-fold and 75-fold differences respectively (Figure 2a). Interestingly, *Pou2f3*, a transcription factor important in differentiation of MVCs (Yamaguchi et al., 2014; Yamashita et al., 2017), is found within the top 10 upregulated genes found in MVC_eGFP cells compared to OSN_eGFP- (Figure 2a). In addition, transcripts for chemosensory cell specific cytokine IL-25 and its receptor IL-17RB (Ualiyeva et al., 2020) were more highly expressed in MVC_eGFP (Figure 2 – figure supplement 1). Finally, *OMP* and *s100a5*, genes for two proteins expressed in mature OSNs (Farbman and Margolis, 1980; Fischl et al., 2014), were among the top 10 downregulated transcripts in MVC_eGFP cells compared to OSN_eGFP- cells (Figure 2a).

**Figure 2.**
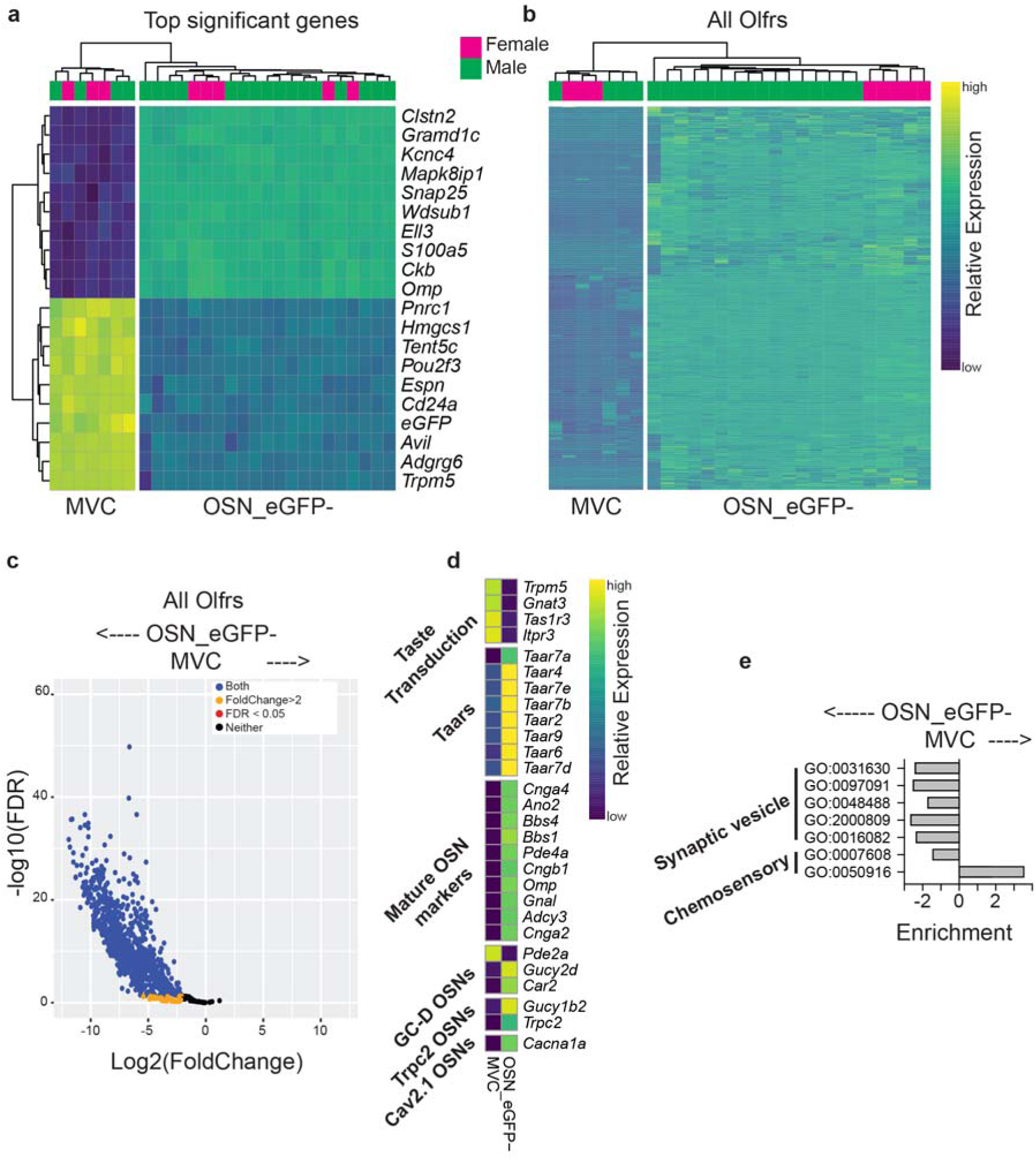
RNAseq comparison of MVC_eGFP vs. OSN_eGFP- cells. **a.** Heatmaps showing hierarchical clustering of the top 10 upregulated and top 10 downregulated genes identified by DESeq2. **b.** Heatmaps showing hierarchical clustering of the 550 olfactory receptor genes identified by DESeq2 as expressed in OSN_eGFP- cells. For both a and b, row and column order were determined automatically by the *pHeatmap* package in R. Row values were centered and scaled using ‘scale = “row”’ within *pHeatmap*. **c.** Volcano plot of all olfactory receptors, demonstrating the large number of enriched olfactory receptors in the OSN_eGFP- population. **d.** Hierarchical clustering of transcripts for taste transduction and transcripts expressed in canonical and non-canonical OSNs identified by RNAseq as significantly different in expression between MVC_eGFP and OSN_eGFP- cells. The non-canonical OSNs considered here included guanilyl-cyclase D (GC-D) OSNs (Juilfs et al., 1997), Trpc2 OSNs (Omura and Mombaerts, 2014), Cav2.1 OSNs (Pyrski et al., 2018), and OSNs expressing trace amine-associated receptors (Taars) (Liberles, 2015). Transcripts identified by DESeq2. **e.** Gene ontology (GO) term enrichment for synaptic vesicle or chemosensory-related GOs was calculated from differentially expressed genes using *TopGO* in R. An enrichment value for genes with Fischer p value <0.05 was calculated by dividing the number of expressed genes within the GO term by the number expected genes (by random sampling, determined by *TopGO*).

We compared expression of transcripts involved in taste transduction, canonical olfactory transduction, and non-canonical OSNs (Figure 2d). MVC_eGFP cells expressed genes involved in the taste transduction pathway as expected for chemosensory epithelial cells of the olfactory epithelium (Ualiyeva et al., 2020). In contrast, OSN_eGFP- expressed transcripts for markers of canonical OSNs such as OMP, BBS1 and 2 and proteins involved in the canonical olfactory transduction pathway. The non-canonical OSNs considered here included guanilyl-cyclase D (GC-D) OSNs (Juilfs et al., 1997), Trpc2 OSNs (Omura and Mombaerts, 2014) and Cav2.1 OSNs (Pyrski et al., 2018). OSN_eGFP- expressed low levels of *Cancna1a* encoding for Cav2.1 and *Trpc2*. OSN_eGFP- expressed higher levels of Trace amine-associated receptors (Liberles, 2015) than MVC_eGFP cells.

Perusal of these top differences suggested that these are distinct chemosensory cell types found in the olfactory epithelium. In order to perform a thorough analysis of the differences between these chemosensory cell groups we performed an analysis of gene ontology (GO) enrichment for lists of genes related to chemosensory perception and neuronal identity. When compared with OSN_eGFP- we found that MVC_eGFP cells were enriched for transcripts belonging to gene ontologies of sensory perception of sweet/umami taste (GO:0050916 and GO:0050917) (Figure 2e, Figure 2 - figure supplement 2, Figure 2 - figure supplement 3) which includes taste detection/transduction proteins that have been reported to be expressed in MVCs (Genovese and Tizzano, 2018; Hegg et al., 2010): *Gnat3*, encoding for gustducin, the G protein mediating sweet and umami taste transduction (McLaughlin et al., 1992), *Itpr3*, encoding for the inositol-1,4,5- triphosphate receptor type 3 and *Tas1r3*, encoding for a gustducin-coupled receptor involved in umami and sweet taste (Damak et al., 2003; Zhang et al., 2003). In contrast, OSN_eGFP- were enriched for transcripts involved in events required for an organism to receive an olfactory stimulus, convert it to a molecular signal, and recognize and characterize the signal (GO:0007608). Finally, enrichment of gene ontology lists for synaptic vesicle function were decreased for MVC_eGFP cells compared with OSN_eGFP- cells (Figure 2e). Results of this gene ontology analysis of chemosensation and synaptic vesicle function reinforces the finding that the two cell groups in this study are distinct chemosensory cell types of the olfactory epithelium. OSN_eGFP- cells differ from MVC_eGFP cells in expression of olfactory receptors, chemosensation and transcripts related to synaptic function as expected for an OSN.

Finally, a question that arises is how the transcriptional profile of the MVC_eGFP cells of this study compares to transcriptional profiling of chemosensory epithelial cells isolated from the respiratory and olfactory epithelia in mice expressing eGFP under control of the ChAT promoter (Ualiyeva et al., 2020). Figure 2 - figure supplement 4 shows comparisons of gene expression between MVC_eGFP and OSN_eGFP- cells of this study and ChAT-eGFP MVCs and ChAT-eGFP SCCs profiled in the respiratory epithelium in the study of Ualiyeva and co-workers (Ualiyeva et al., 2020). This comparison is of limited value due to the fact that gene profiling was performed in two separate studies. However, in this preliminary analysis we find that MVCs from this study have similar transcription profiles to ChAT-eGFP MVCs and differ from ChAT-eGFP SCCs. For example, MVC_eGFP and ChAT-eGFP MVCs showed enhanced expression of transcripts such as *Il25* and *Fos* (Figure 2 - figure supplement 3). The similarity of transcriptional profiling argues that in this study MVC_eGFPs were not contaminated with SCCs consistent with the fact that in our study we isolated MVC_eGFP from olfactory epithelium dissected apart from the respiratory epithelium and that the density of MVCs in the OE is higher than the density of SCCs in the respiratory epithelium (Ualiyeva et al., 2020) decreasing the chance of contamination of OE MVCs by SCCs. Interestingly, *Ugt2a1* and *Ugt2a2*, transcripts for proteins involved in UDP synthesis were higher in MVC_eGFP than ChAT-eGFP MVCs suggesting differences between these cells (Figure 2 - figure supplement 3). In order to determine whether these similarities and differences between MVCs in our study and the study of Ualiyeva and co-workers are real it will be necessary to perform simultaneous RNAseq profiling of these two populations.

### Gene ontology analysis finds enrichment of lists of viral-related, inflammation and immune transcripts in MVC_eGFP cells

SCCs, tuft and brush cells have been implicated in responses to bacterial and viral infection, immunity and inflammation (Luo et al., 2019; Maina et al., 2018; O’Leary et al., 2019; Perniss et al., 2020; Rane et al., 2019; Saunders et al., 2014; Tizzano et al., 2010; Ualiyeva et al., 2020). The fact that MVCs are closely related to these cells (Fu et al., 2018; Genovese and Tizzano, 2018; Ogura et al., 2011; Ualiyeva et al., 2020) lead us to search for gene ontology enrichment related to bacterial and viral infection, immunity and inflammation for MVC_eGFP cells. We found robust enrichment of these gene ontologies in MVC_eGFP cells (Figure 3a). Transcripts related to viral infection that were higher in MVC_eGFP cells compared to OSN_eGFP- cells (Figure 3b) included those involved in viral entry into host cells, viral transcription and regulation of viral transcription, negative regulation of viral genome replication and negative regulation of viral process (Figure 2 – figure supplement 2). The majority of these genes were detected by Ualiyeva and colleagues (Ualiyeva et al., 2020) in their ChAT-GFP MVC population. We also found gene ontology enrichment in MVC_eGFP cells compared to OSN_eGFP- cells for defense response to bacterium (Figure 2 – figure supplement 2).

**Figure 3.**
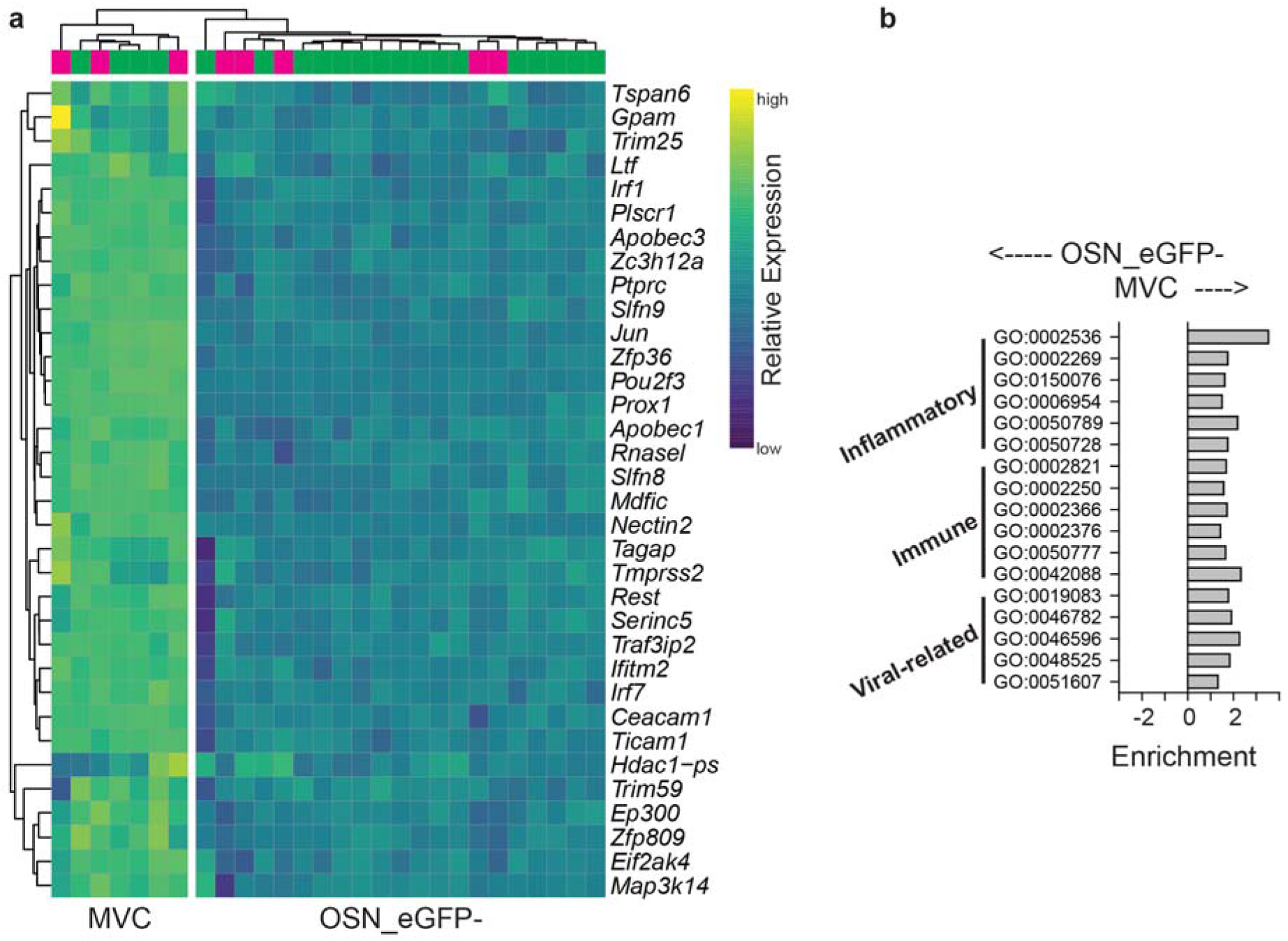
Significant differences in virally-related, immune and inflammation gene ontology lists between MVC_eGFP and OSN_eGFP-. **a.** Gene ontology (GO) term enrichment was calculated from differentially expressed genes using *TopGO* in R for OSN_eGFP- vs. MVC_eGFP cells. An enrichment value for genes with Fischer p value <0.05 was calculated by dividing the number of expressed genes within the GO term by the number expected genes (by random sampling, determined by *TopGO*). Heatmap show hierarchical clustering of significantly differentially expressed genes identified by DESeq2. **b.** Significantly differences in virally-related genes within the MVC_eGFP cells compared to OSN_eGFP-.

Importantly, we also find enrichment for transcript expression for immunity and inflammation. Genes related to inflammation and immunity that were higher in MVC_eGFP cells compared to OSN_eGFP- cells are shown in Figure 3 – figure supplements 1-2. Among these transcripts *IL25* and its receptor *Il17rb* are enriched in MVC_eGFP cells. In SCCs, brush cells and tuft cell generation of IL25 leads to a type 2 inflammation and stimulates chemosensory cell expansion in a sequence of events that also involves cysteinyl leukotrienes (Bankova et al., 2018; Luo et al., 2019; von Moltke et al., 2016). The presence of both *Il25* and *Il17rb* suggests an autocrine effect. Furthermore, both cell types displayed increased expression of transcripts encoding for enzymes involved in eicosanoid biosynthesis such as *Alox5*, *Ptgs1* and *Ptgs2* that are found in brush cells in the airways (Bankova et al., 2018) and tuft cells in the intestine (McGinty et al., 2020) where they drive type 2 immune responses.

### Transcription profiling indicates that OSN_eGFP+ cells are distinct from both OSN_eGFP- and MVC_eGFP cells

Differential gene expression analysis of the RNAseq data was used to compare OSN_eGFP+ individually with the other two groups of cells. We found that expression of 2000 genes was significantly higher in OSN_eGFP+ compared to OSN_eGFP-, and expression of 1821 genes was lower in OSN_eGFP+ cells (Figure 4 -figure supplement 1 shows the results of RNAseq and Figure 4 -figure supplement 2 summarizes the data). Figure 4 figure supplement 2a shows expression levels for the transcripts that showed the largest differences between OSN_eGFP+ and OSN_eGFP- cells. The transcripts for TRPM5 and eGFP were among the top 10 genes whose transcription was higher in OSN_eGFP+ compared to OSN_eGFP- with 105-fold and 42-fold increases respectively. However all of these 10 top genes, and many other genes that were found at significantly higher levels of expression in OSN_eGFP+ cells compared to OSN_eGFP- happen to be genes expressed at significantly higher levels in MVC_eGFP cells (Figure 4 -figure supplement 3 shows the results of RNAseq for MVC_eGFP vs OSN_eGFP+). For example *Trpm5* is expressed at levels of 87.5, 9200 and 127000 in OSN_eGFP-, OSN_eGFP+ and MVC_eGFP cells respectively (Figure 4 -figure supplement 4). While the light scatter settings in the FACS were set to exclude doublets, this raised the question whether expression of these genes in the OSN_eGFP+ pool was due to contamination of the OSN_eGFP+ cell fraction (mCherry and eGFP positive) by doublets made up of one OSN_eGFP- cell (mCherry positive and eGFP negative) and one MVC_eGFP cell (mCherry negative and GFP positive).

In order to determine whether transcription profiling for the OSN_eGFP+ cell fraction is consistent with this being a separate population we searched for genes whose expression levels were significantly higher in OSN_eGFP+ compared to *both* OSN_eGFP- and MVC_eGFP. Figures 4a and 4b show the top genes that were expressed at significantly higher levels in OSN_eGFP+ (and Figure 4 – figure supplement 5 shows data for all 80 genes). Among these genes there were 22 olfactory receptor genes and one olfactory receptor pseudogene (Figure 4b, Figure 4c shows a volcano plot for Olfrs). A GOnet GO term enrichment analysis (Pomaznoy et al., 2018) of the 80 genes enriched in OSN_eGFP+ compared to the other two groups (Figure 4 – figure supplement 6) revealed that these cells express genes involved in sensory perception of smell (GO:0007608), signal transduction (GO:0007165) and cellular response to stimulus (GO:0051716). Interestingly, two of these genes *Trpc2* (Omura and Mombaerts, 2014) and *Calb2* (Bastianelli et al., 1995) are expressed in small subsets of OSNs. Thus, this analysis indicates that OSN_eGFP+ cells are distinct from the other two cell populations, although more detailed follow-up experiments are necessary to fully characterize this population including single cell RNA sequencing.

**Figure 4.**
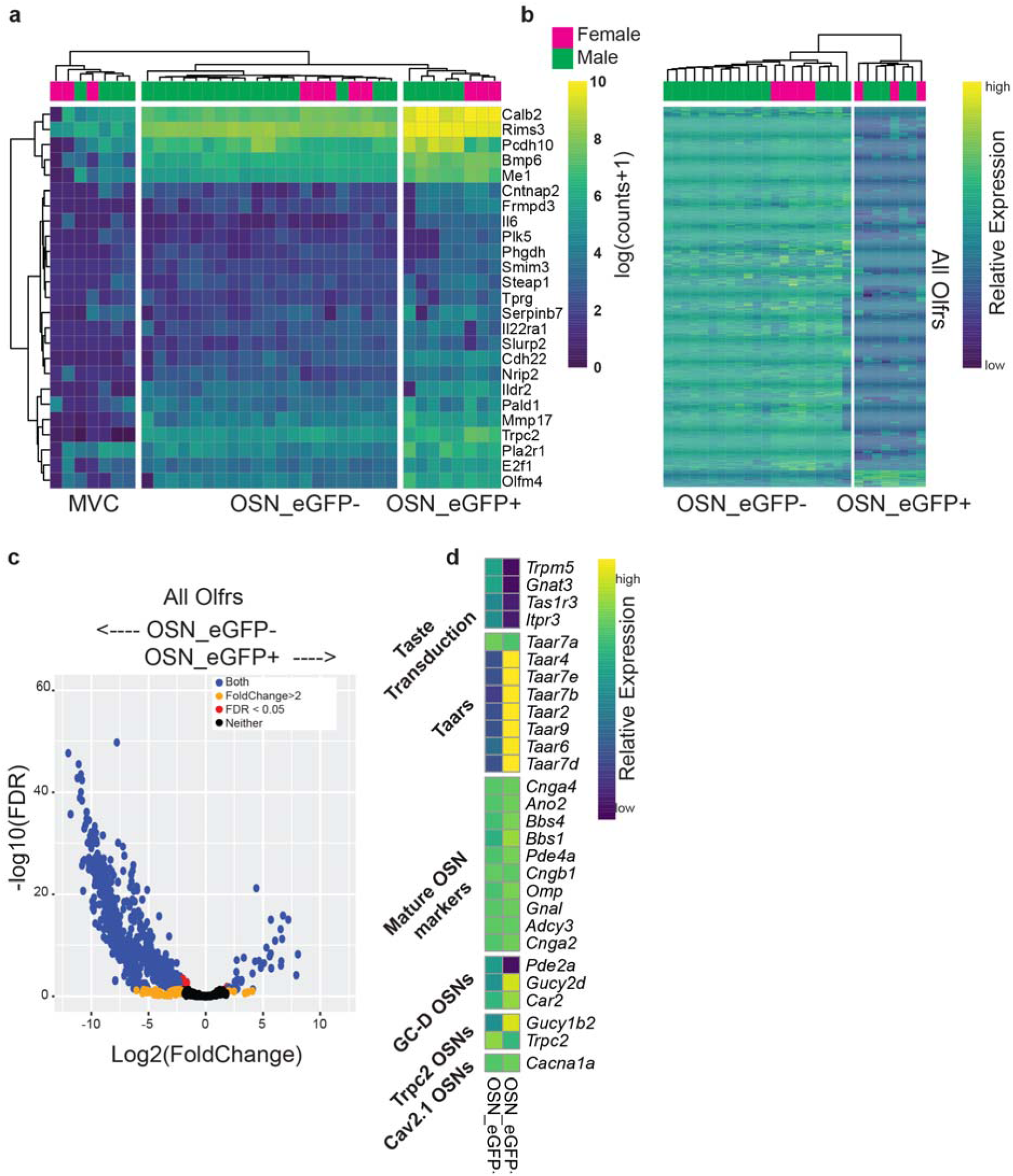
RNAseq comparison of OSN_eGFP+ to both MVC_eGFP and OSN_eGFP- cells. **a.** Heatmap showing the top upregulated genes (excluding Olfrs) that are expressed in OSN_eGFP+ cells 4 fold higher than OSN_eGFP- AND MVC_eGFP cells. Additional criteria for inclusion was mean of expression > standard deviation of expression and mean of expression greater than 100. **b.** Heatmap showing all *Olfr* genes differentially expressed between OSN_eGFP+ and OSN_eGFP- cells identified by DESeq2. MVC_eGFP cells did not express Olfrs. For both a and b, row and column order were determined automatically by the *pHeatmap* package in R. For each data point relative expression was calculated by subtracting the average row value from each individual value. **c.** Volcano plot of all Olfactory receptors, demonstrating the small number of enriched olfactory receptors in the OSN_eGFP+ population. **d.** Hierarchical clustering of transcripts for taste transduction and transcripts expressed in canonical and non-canonical OSNs identified by RNAseq as significantly different in expression between the cell groups. We compared expression of transcripts involved in taste transduction, canonical olfactory transduction, and non-canonical OSNs. The non-canonical OSNs considered here included guanilyl-cyclase D (GC-D) OSNs (Juilfs et al., 1997), Trpc2 OSNs (Omura and Mombaerts, 2014), Cav2.1 OSNs (Pyrski et al., 2018), and OSNs expressing trace amine-associated receptors (Taars) (Liberles, 2015). Transcripts identified by DESeq2.

### Gender differences for expression of olfactory receptors

We did not find major differences in transcriptome profiling between males and females for genes that were differentially expressed between the three cell groups (Figure 4 – figure supplement 7,8). We found a substantial number of olfactory receptor genes that were differentially expressed between males and females (Figure 4 – figure supplement 8). Interestingly, *Trpc2*, that is one of the genes with higher expression in OSN_eGFP+ cells is expressed in higher amounts in females. Surprisingly, the differentially expressed olfactory receptors differed from receptors identified by van der Linden et al. (van der Linden et al., 2018).

### *In situ* hybridization chain reaction finds strong TRPM5 mRNA expression in MVC_eGFP cells, but not in the nuclear OSN layer

Studies with regular *in situ* hybridization find expression of TRPM5 mRNA in MVCs, but not in the OSN nuclear layer (Pyrski et al., 2017; Yamaguchi et al., 2014). Here we asked whether third generation in situ hybridization chain reaction version 3.0 (HCR v3.0) designed to provide high signal to noise ratio *in situ* signal (Choi et al., 2018) revealed TRPM5 mRNA expression in the nuclear OSN layer. These experiments were performed in TRPM5-GFP mice and in TRPM5-GFP mice crossed with TRPM5 knockouts (Clapp et al., 2006; Damak et al., 2006). Consistent with published results (Pyrski et al., 2017; Yamaguchi et al., 2014) we find strong *in situ* signal for TRPM5 in MVC_eGFP cells located in the apical layer of the olfactory epithelium (Figure 5a, asterisks, also see Figure 5 – figure supplement 1 for a 3D rendering) and this signal is absent in MVC_eGFP cells in the TRPM5 knockout (Figure 5b, asterisks). In addition, we find sparse TRPM5 *in situ* labeling in the nuclear OSN layer (Figure 5a, arrows), but similar sparse labeling was found in the OSN nuclear layer in the TRPM5 knockout (Figure 5b, arrows). Therefore, we find evidence for strong expression of TRPM5 mRNA in MVC_eGFP cells, but we do not find evidence for expression of TRPM5 mRNA in OSNs.

**Figure 5.**
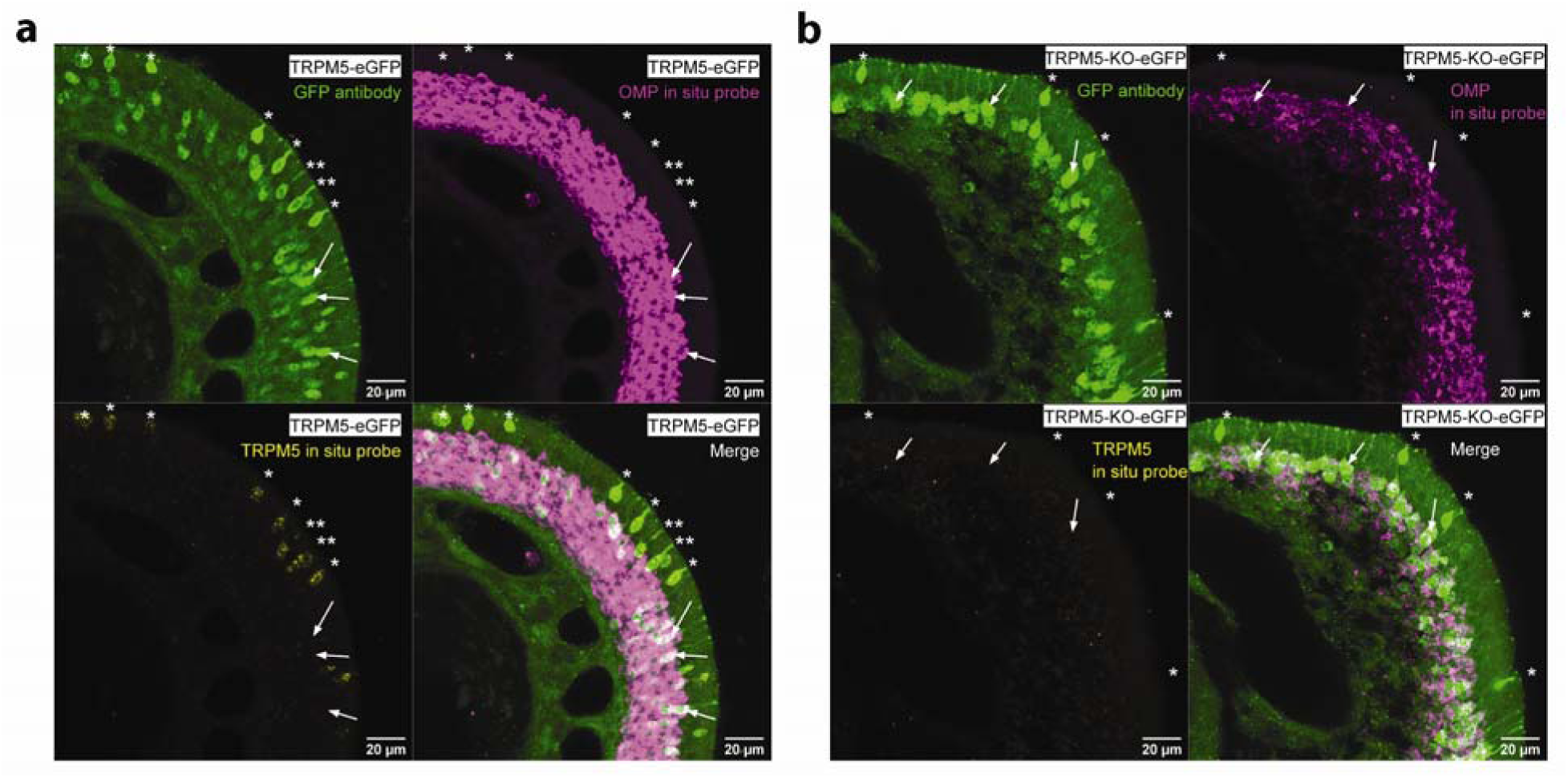
*In situ* hybridization chain reaction finds strong TRPM5 mRNA expression in MVC_eGFP cells, but not in the nuclear OSN layer. **a.** In situ for TRPM5 (yellow) and OMP (magenta) transcripts in the olfactory epithelium of TRPM5-GFP mice (GFP is green) shows strong label for TRPM5 in MVCs (asterisks) and sparse labeling in the OSN nuclear layer (arrows). The scale bar is 20 μm. **b.** In situ for TRPM5 (yellow) and OMP (magenta) transcripts in the olfactory epithelium of TRPM5-GFP x TRPM5-knockout mice (GFP is green) shows no label for TRPM5 in GFP- positive MVCs (asterisks) and does show sparse labeling in the OSN nuclear layer (arrows). The scale bar is 20 μm.

## Discussion

We performed transcriptional profiling of three chemosensory cells in the mouse olfactory epithelium: microvillous cells (MVC_eGFP) and two types of olfactory sensory neurons: OSN_eGFP+ and OSN_eGFP-. We found that while the transcriptome of each of these cell types is distinct they share common features across groups. The two groups of OSNs share transcript expression for proteins expressed in OSNs such as OMP, olfactory transduction proteins, and proteins involved in synaptic function. Yet, they differ in olfactory receptor expression and OSN_eGFP+ express transcripts encoding for proteins involved in chemosensory signal transduction and cellular response to stimulus. On the other hand, MVC_eGFP cells express transcripts encoding for taste transduction proteins and other transcripts found in SCCs such as *Pou2f3* and *Il25* but they do not express transcripts for proteins involved in olfactory transduction and synaptic function, and they do not express olfactory receptors. Finally, we found that MVC_eGFP cells express a substantial number of transcripts involved in viral infection, inflammation and immunity.

### Transcriptional profiling reveals a role of microvillous cells in viral infection and innate immunity

Gene ontology analysis revealed that MVC_eGFP cells are enriched in viral-related transcripts compared to OSN_eGFP- (Figure 3 and Figure 2 – figure supplement 2). GOnet GO term enrichment analysis (Pomaznoy et al., 2018) of all 133 immune genes enriched in MVC_eGFP cells compared to OSN_eGFP- (Figure 3 – figure supplement 2) revealed that MVCs express a substantial number of genes involved in the innate immune response (GO:0045087, 72 genes matched this list). Figure 6 depicts several mechanisms that could occur in MVC_eGFP in response to viral infection. To infect cells, viruses must interact with host cell membranes to trigger membrane fusion and viral entry. Membrane proteins at the surface of the host cell are thus key elements promoting or preventing viral infection. Here we find that transcripts for several membrane proteins and cell adhesion molecules involved in viral entry are enriched in MVC_eGFP cells. *Plscr1* encodes a phospholipid scramblase which has been shown to promote herpes simplex virus (HSV) entry in human cervical or vaginal epithelial cells and keratinocytes (Cheshenko et al., 2018), and hepatitis C virus entry into hepatocytes (Gong et al., 2011). In contrast with its role in viral entry, PLSCR1 impairs the replication of other types of viruses in infected cells (influenza A virus (Luo et al., 2018), hepatitis B virus (Yang et al., 2012)). IFTM2 is another transmembrane protein that mediates viral entry. In contrast with PLSCR1, IFTM2 inhibits viral entry of human immunodeficiency virus (HIV, (Yu et al., 2015)), hepatitis C virus (Narayana et al., 2015), influenza A H1N1 virus, West Nile virus, and dengue virus (Brass et al., 2009). IFTM2 also inhibits viral replication (Brass et al., 2009) and protein synthesis (Lee et al., 2018). Nectins are transmembrane glycoproteins and constitute cell surface receptors for numerous viruses. There is wide evidence that HSV can enter host cells through Nectin-1 dependent mechanisms, particularly for neuronal entry (Kopp et al., 2009; Petermann et al., 2015; Sayers and Elliott, 2016; Shukla et al., 2012), and Nectin-4 appears essential for measles virus epithelial entry (Noyce and Richardson, 2012; Singh et al., 2015; Singh et al., 2016). In addition to cell surface molecules, the mucus contains secreted proteins that confer protection against viruses to the underlying cells. Glycoproteins are major constituents of mucus and exhibit multiple pathogens binding-sites. We found the *Ltf* transcript in MVC_eGFP cells, which encodes for lactotransferrin. Lactotransferrin is a globular glycoprotein widely represented in the nasal mucus with anti-viral activity against Epstein-Barr virus (Zheng et al., 2014; Zheng et al., 2012), HSV (Shestakov et al., 2012; Valimaa et al., 2009)) and Hepatitis C virus (Allaire et al., 2015). Finally, MVC_eGFP cells express the murine norovirus (MNoV) receptor CD300LF. In the gut TRPM5-expressing tuft cells express high levels of CD300LF and mice were resistant to infection with MNoV^CR6^ when tuft cells were absent or decreased, whereas viral titers were enhanced in any context where tuft cell numbers were increased, such as helminth infection or treatment with rIL-25 (Wilen et al., 2018).

**Figure 6.**
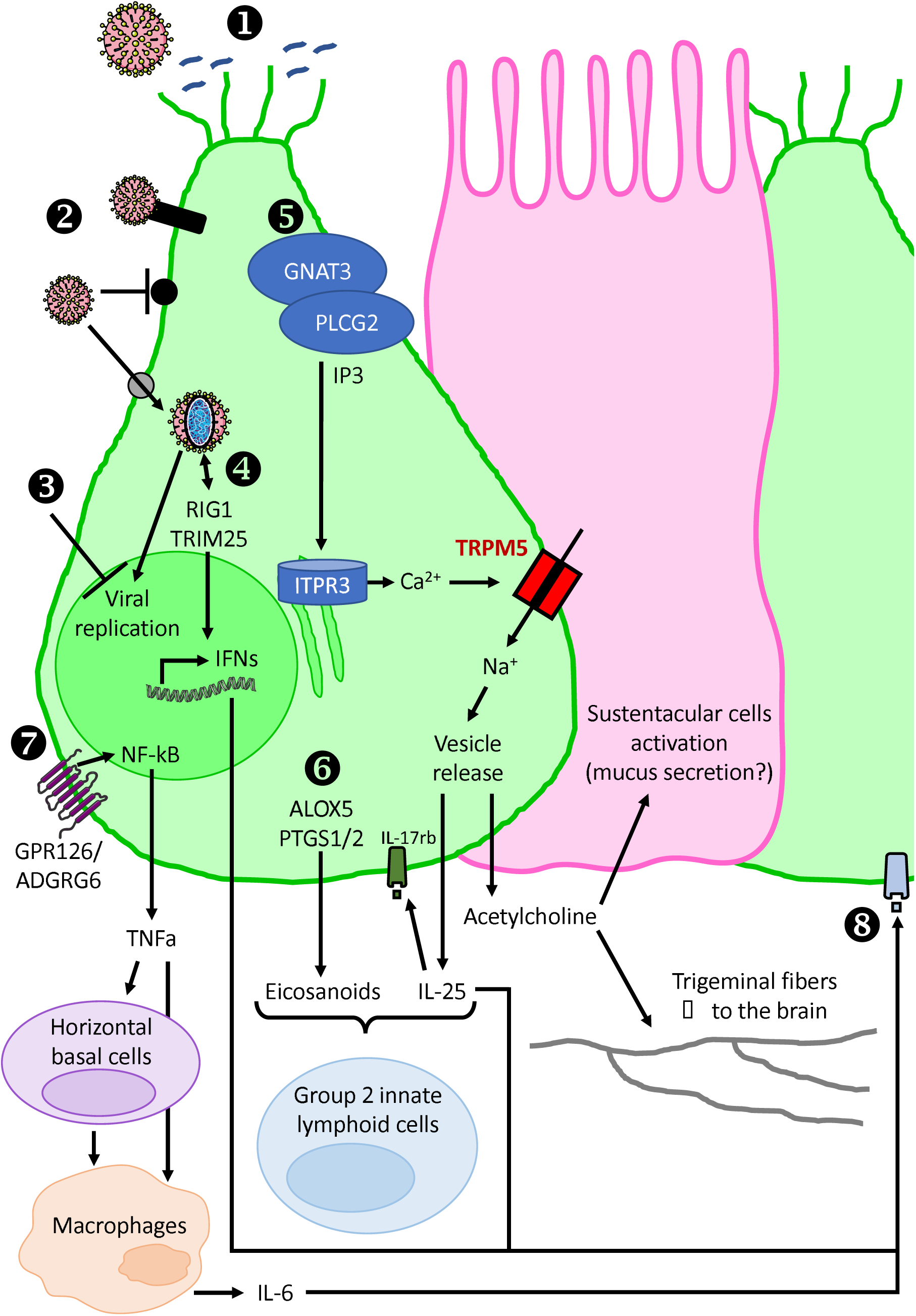
Model depicting the role for microvillous cells involvement in the olfactory epithelium innate immune response to viral infection. 1) Secreted or cell surface glycoproteins constitute a first barrier preventing virus entry. 2) When reaching MVC_eGFP, viruses can encounter three types of membrane proteins: adhesion molecules that trigger intracellular signaling upon viral recognition (black rectangle), transmembrane proteins that block virus entry (black circle), viral receptors allowing virus entry (grey circle). 3) MVC_eGFP express numerous transcriptional factors involved in the inhibition of viral replication. 4) Cytosolic viral RNA sensing induces the production of type I interferons. 5) A possible signaling pathway leading to intracellular calcium increase, TRPM5 activation and Na^+^-mediated vesicle release. Acetylcholine can activate neighboring sustentacular cells and underlying trigeminal fibers. 6) Eicosanoids synthesis, along with IL-25 production, can recruit and activate group 2 innate lymphoid cells, which are key controllers of type 2 inflammation. 7) GPR126 activation results in NFkB activation and TNFα production. TNFα can directly activate macrophages. TNFα also induces a change in the function of horizontal basal cells, switching their phenotype from neuroregeneration to immune defense. 8) Interferons and cytokines can in turn activate antiviral immune response in neighboring MVC_eGFP.

Viruses have developed numerous strategies to overcome barrier mechanisms to enter the cells. After viral entry infected cells have other resources to fight against viral infection by disrupting the production of new viral particles, limiting inflammation processes and activating innate immune responses. For example, TRIM25, whose transcript is increased in MVC_eGFP cells, is an ubiquitin ligase that activates retinoic acid-inducible gene I (RIG-I) to promote the antiviral interferon response (Gack et al., 2007). Furthermore, influenza A virus targets TRIM25 to evade recognition by the host cell (Gack et al., 2009). In addition, TRIM25 displays a nuclear role in restricting influenza A virus replication (Meyerson et al., 2017). Zc3h12a, also known as MCPIP-1, inhibits hepatitis B and C virus replication, reduces virus-induced inflammation (Li et al., 2020; Lin et al., 2014), and exerts antiviral effects against influenza A virus (Dong et al., 2017).

We show that MVC_eGFP express *Gnat3*, *Plcg2* and *Itpr3*. This intracellular pathway would lead to calcium increase and opening of TRPM5, leading to a sodium influx and potential vesicle release. Among the list of inflammation genes enriched in MVC_eGFP we find *Il25*, an interleukin that is involved in the type 2 inflammatory response of TRPM5-expressing epithelial cells in the airway epithelium and the gut (O’Leary et al., 2019; Ting and von Moltke, 2019). Also, *Il25* expression in the skin leads to disruption of the epithelium and enhances HSV-1 and vaccinia virus replication (Kim et al., 2013). MVC_eGFP cells are known to produce acetylcholine, which can activate sustentacular cells through M3 muscarinic acetylcholine receptors (Ogura et al., 2011). Sustentacular cells may in particular play a role in maintaining extracellular ionic gradients, extracellular glucose, secreting mucus, metabolizing noxious chemicals, and regulating cell turnover (Fu et al., 2018; Villar et al., 2017). In addition to *Il-25*, the expression of enzymes for eicosanoid biosynthesis (*Alox5*, *Ptgs1* and *Ptgs2*) suggests that MVC_eGFP are likely to recruit group 2 innate lymphoid cells, similar to tuft cells in the small intestine (McGinty et al., 2020). Finally, the innate immune response involves recruitment of macrophages that are known to play a protective role in the olfactory epithelium (Borders et al., 2007). MVC_eGFP express the G-protein coupled receptor GPR126/ADGRG6, which is required for macrophage recruitment and Schwann cells regeneration after peripheral nerve injury (Mogha et al., 2016). This raises the question whether MVC_eGFP could play a protective role, promoting OSN survival and increasing neurogenesis, through macrophage recruitment. In addition, activation of MVCs by irritants, bacteria and viruses could result in activation of cytokine-induced inflammation and macrophage recruitment by long-term horizontal basal cells, that activate type 1 immune responses within the olfactory epithelium (Chen et al., 2019). All cytokines and interferons produced in the microenvironment of a MVC_eGFP can then contribute to the activation of immune responses in neighboring MVC_eGFP, since we found the expression of various cytokine receptors (*IL6ra, Il1rap, IL4ra, IL17re, IL17 rb, TNFRSF13B*) and interferons responsive elements (*Ifitms*).

Our findings of expression of virally relevant transcripts in MVC_eGFP cells complement published studies on the role of MVC-related SCCs in viral infection. In the trachea, viral-associated formyl peptides activate SCCs to release acetylcholine and activate mucocilliary clearance by ciliated cells (Perniss et al., 2020). This activation is mediated by the TRPM5 transduction pathway in the SCC and muscarinic acetylcholine receptors in the ciliated cell. In a similar manner in the olfactory epithelium MVCs respond to ATP, which is involved in activating mucociliary movement by releasing acetylcholine and activating adjacent sustentacular cells through a muscarinic receptor (Fu et al., 2018). Therefore, viral infection could result in activation of MVCs resulting in activation of mucociliary clearance by adjacent sustentacular cells.

*Pou2f3* also called *Skn1a*, encodes for a key regulator for the generation of TRPM5-expressing cells in various epithelial tissues (Yamashita et al., 2017). *Pou2f3* transcript was increased in MVC_eGFP cells compared to OSN_eGFP-. *Skn1a/Pou2f3*-deficient mice lack intestinal tuft cells and have defective mucosal type 2 responses to helminth infection in the intestine (Gerbe et al., 2016). MVCs express markers of tuft cells (*Pou2f3*, *Trpm5* and others) indicating that MVCs share inflammatory and innate immune functions with tuft cells in the gut and SCCs and brush cells in the airways (Ting and von Moltke, 2019). In addition, in the anterior olfactory epithelium, where there is a higher density of MVCs, mice exposed to mild odorous irritants exhibited a time-dependent increase in apoptosis and a loss of mature OSNs without a significant increase in proliferation or neurogenesis (Lemons et al., 2020). Future experiments are necessary to determine whether activation of MVCs by viruses could lead to loss of mature OSNs contributing to smell loss after viral infection. Interestingly, in the mouse distal lung, where there is no expression of SCCs, there was de novo generation of SCCs after infection with A/H1N1/PR/8 influenza virus (Rane et al., 2019) raising the question whether virus exposure could alter MVC number in the olfactory epithelium. Finally, in teleost fish rhabdoviruses induce apoptosis in a unique type of crypt OSN via the interaction of the OSN TrkA receptor with the viral glycoprotein and activates proinflammatory responses in the olfactory organ (Sepahi et al., 2019).

### Viral infection of the central nervous system through the olfactory epithelium

The olfactory epithelium provides direct viral access to the brain through the olfactory nerve. Whether this olfactory path constitutes route of entry for viruses to the brain is a matter of intense discussion, especially because some viruses are postulated to be involved in encephalopathy and neurodegenerative disorders (Cairns et al., 2020; Dando et al., 2014; Doty, 2008; Harris and Harris, 2018). Our finding that MVC_eGFP cells are enriched in virally-related genes suggests that these cells may be involved in or prevention of viral entry into the brain (and these two alternatives are not exclusive since they may be different for different viruses). On the one hand, we identified transcripts encoding for viral receptors in MVC_eGFP, suggesting that viruses can enter these cells. It is not known whether viruses can spread in neighboring cells, but if viral particles were to enter the OSNs they could reach the olfactory bulb through anterograde transport along the olfactory nerve and from the olfactory bulb, viruses can spread throughout the brain along the olfactory bulb-hippocampus route. On the other hand, we found enrichment for transcripts encoding for proteins involved in limiting viral infection and promoting immune and anti-inflammatory responses in MVC_eGFP cells. In this case, viral spread to the brain would be prevented. Finally, the olfactory epithelium is innervated by the trigeminal nerve, and substance P immunostaining is closely associated with subsets of MVCs (Lin et al., 2008). This raises the question whether an interaction between MVCs and trigeminal nerve fibers could participate in local inflammation as found for SCCs (Saunders et al., 2014), and could modulate the entry of virus to the brain stem through the trigeminal nerve. Future experiments are necessary to study the potential role of MVC_eGFP cells in viral infection of the olfactory epithelium and the brain.

### Are GFP-expressing OSNs in the TRPM5-GFP mouse a distinct set of OSNs?

Expression of TRPM5 in a subset of OSNs has been controversial. The original proposal of expression of TRPM5 in OSNs was motivated by expression of eGFP in adult TRPM5-eGFP transgenic mice and immunolabeling of the ciliary layer of the epithelium with an antibody (raised against the TRPM5 peptide RKEAQHKRQHLERDLPDPLDQK) that was validated by lack of expression in TRPM5 knockout mice(Lin et al., 2007). However, staining in the ciliary layer with this antibody has not been replicated and our group and others subsequently showed knockout-validated TRPM5 staining of microvillous cells, and no labeling of the ciliary layer in the adult with antibodies raised against different TRPM5 protein peptides(Genovese and Tizzano, 2018; Gilbert et al., 2015; Lin et al., 2008; Pyrski et al., 2017). Interestingly, Pyrski and co-workers found TRPM5 immunolabeling in OSNs in the embryo, but not in the adult(Pyrski et al., 2017). Furthermore, in the adult mouse *in situ* hybridization has reported mRNA staining for TRPM5 in MVCs, and not in OSNs(Pyrski et al., 2017; Yamaguchi et al., 2014), and full length TRPM5 mRNA was not found in OSNs in the adult(Pyrski et al., 2017).

Here we corroborate expression of eGFP in OSNs in TRPM5-eGFP transgenic mice. In addition, using HCR v3.0 we find strong expression of mRNA in MVCs, but we do not find evidence for expression of TRPM5 mRNA in OSNs (Figure 5). However, transcriptional analysis indicates that the OSN_eGFP+ cell population differs in mRNA expression from the other two populations (Figure 4). OSN_eGFP+ express a subset of olfactory receptors as well as transcripts encoding for proteins involved in chemosensory signal transduction and cellular response to stimulus. This is consistent with calcium imaging and loose patch recordings from adult TRPM5-eGFP transgenic mice that found that GFP-labeled cells with morphology resembling OSNs (presumably OSN_eGFP+ cells) responded to pheromones and MHC peptides with currents that were abolished by pharmacologic inhibition of TRPM5 or isolation of cells from TRPM5 knockouts(Lopez et al., 2014). In addition, studies of neuronal connectivity found that approximately half of the glomeruli innervated by GFP-bearing axons in the adult TRPM5-eGFP transgenic mouse are innervated by mitral cells that project directly to the medial amygdala, consistent with these glomeruli carrying pheromone information(Thompson et al., 2012). Nevertheless, these findings are in contrast with findings by Pyrksi and co-workers who show that in adult Trpm5-IRES-Cre x R26-GCaMP3 mouse OSNs that express GCaMP3 do not respond selectively to pheromones, and responded to general odorants(Pyrski et al., 2017). Why the functional studies of Lopez and co-workers and Pyrski and co-workers differ is not clear. However, eGFP-bearing OSNs in the Trpm5-IRES-Cre crossed with an eGFP reporter were expressed throughout the olfactory epithelium with no obvious spatial pattern (Pyrski et al., 2017) in contrast with the expression of eGFP in TRPM5-eGFP transgenics that is stronger in the lateral olfactory epithelium(Oshimoto et al., 2013). This raises the question whether eGFP is expressed in different OSNs in the Trpm5-IRES-Cre and TRPM5-eGFP transgenics. In summary, our data indicate that OSN_eGFP+ cells are a distinct population of chemosensory cells, but whether these are pheromone-responding OSNs requires future studies.

### Would microvillous cells play a role in COVID-19?

Recently, due to the current COVID-19 pandemic, researchers have focused their attention on investigating SARS-CoV-2 mechanism of entry into cells. SARS-CoV-2 targets mainly cells of the respiratory pathway where viral entry is mediated by ACE2 and TMPRSS2 (Hoffmann et al., 2020). Because numerous patients reported loss of smell (Giacomelli et al., 2020; Parma et al., 2020; Yan et al., 2020a; Yan et al., 2020b), researchers wondered about the mechanism for SARS-CoV-2 infection of the olfactory epithelium. In our study, we found the *Tmprss2* transcript was significantly increased in MVC_eGFP cells compared to OSN_eGFP- (Figure 3). We did not find *Ace2* enrichment in these cells, but this may be due to inefficiency in finding with RNAseq low abundance transcripts like *Ace2* (Ziegler et al., 2020). Transcriptional profiling of single cells in the olfactory epithelium from other laboratories found expression of transcripts for both *Tmprss2* and *Ace2* in in sustentacular cells and stem cells, and at lower levels in MVCs (Brann et al., 2020; Fodoulian et al., 2020). Viral infection of sustentacular cells may explain loss of smell because these cells play a key role in supporting olfactory function by providing glucose for the energy necessary for olfactory transduction in the OSN cilia (Villar et al., 2017). Importantly, type I interferons, and to a lesser extent type II interferons induced by response of the host to SARS-CoV-2, and infection by other viruses inducing the interferon pathway increases *Ace2* expression in the nasal epithelium (Ziegler et al., 2020). MVCs may play a role in SARS-CoV-2 infection of the olfactory epithelium because these cells may participate in activating inflammation of the epithelium that elicits type 1 immune response (Chen et al., 2019).

### Conclusion

Here we find that microvillous cells of the olfactory epithelium express transcripts involved in immunity, inflammation and viral infection. These expression patterns suggest that, like tuft cells in the gut and SCCs and brush cells in the airways, the microvillous cells in the olfactory epithelium are involved in the innate immune response to viral infection. Our study provides new insights into a potential role for TRPM5-expressing cells in viral infection of the main olfactory epithelium.

## Abbreviations

COVID-19: Coronavirus disease 2019
DPI: Days post infection
eGFP: Enhanced green fluorescent protein
FACS: Fluorescence-activated cell sorting
FDR: False discovery rate
GLM: Generalized linear model
GO: Gene ontology
MVCs: Microvillous cells
OMP: Olfactory marker protein
OSNs: Olfactory sensory neurons
PSF: Point spread function
SARS-CoV-2: Severe acute respiratory syndrome coronavirus clade 2
SSC: Saline-sodium citrate
SSCT: SSC with tween
TRPM5: Transient receptor potential cation channel subfamily M member 5

## Methods

### Key Resources Table

| REAGENT<br>TYPE | REAGENT or<br>RESOURCE | SOURCE | IDENTIFIER | ADDITIONAL<br>INFORMATION |
| --- | --- | --- | --- | --- |
| Chemical compound, drug | BrainPhys Neuronal Medium | Stemcell Technologies |  | Product # 05791 |
| Chemical compound, drug | Dispase II | Sigma |  | Product # D4693 |
| Chemical compound, drug | AcGFP1/eGFP calibration beads | Takara |  | Flow cytometry calibration beads<br>Product # 632594 |
| Chemical compound, drug | mCherry calibration beads | Takara |  | Flow cytometry calibration beads<br>Product # 632595 |
| Chemical compound, drug | RQ1 RNase-free DNase | Promega |  | Product # M6101 |
| Chemical compound, drug | Papain | Sigma |  | Product # P3125 |
| Chemical compound, drug | Paraformaldehyde (32%) | Electron Microscopy Sciences |  | Product # 157145 |
| Chemical compound, drug | RNAprotect Tissue Reagent | Qiagen |  | Product # 76526 |
| Chemical compound, drug | RNeasy Plus Micro Kit | Qiagen |  | Product # 74034 |
| Chemical compound, drug | High Capacity c-DNA Reverse Transcription kit | ABI |  |  |
| Chemical compound, drug | 18s rRNA | PE ABI |  |  |
| Strain, strain background | TRPM5-eGFP | Dr. Robert Margolskee (Clapp et al., 2006) |  |  |
| Strain, strain background | TRPM5 knockout | Dr. Robert Margolskee {Damak, 2006 #1567} |  |  |
| Strain, strain background | OMP-H2B::Cherry | Generated for this publication |  | This mouse will be deposited in Jackson Laboratories |
| Software, algorithm | MATLAB_R2018a | Mathworks | RRID: SCR_001622 |  |
| Software, algorithm | Illustrator | Adobe | RRID: SCR_010279 |  |
| Software, algorithm | Photoshop | Adobe | RRID: SCR_014199 |  |
| Software, algorithm | InDesign | Adobe |  |  |
| Software, algorithm | MoFlo Astrios Summit Software (6.3.1.16945). | Beckman Coulter |  |  |
| Software, algorithm | BBMap (BBDuk) |  | RRID:SCR_016968 |  |
| Software, algorithm | Salmon v1.2.1 | https://combine-lab.github.io/salmon/ | RRID:SCR_017036 | (Patro et al., 2017) |
| Software, algorithm | DeSEQ2 v1.28.0 | [bioconductor.org<br>https://bioconductor.org/packages/release/bioc/html/DESeq2.html](https://bioconductor.org/packages/release/bioc/html/DESeq2.html) | RRID:SCR_015687 | (Love et al., 2014) |
| Software, algorithm | TopGO, v2.40.0 |  | RRID:SCR_014798 |  |
| Software, algorithm | pHeatmap, 1.0.12 |  | RRID:SCR_016418 |  |
| Software, algorithm | Ensembl GRCm38, v99 |  |  |  |
| Software, algorithm | R, v4.0 |  | RRID:SCR_001905 |  |
| Software, algorithm | Tximport, v1.16.0 |  | RRID:SCR_016752 |  |
| Software, algorithm | SAMtools | SAMtools<br>http://samtools.sourceforge.net/ | RRID:SCR_002105 | (Li et al., 2009) |
| Software, algorithm | Bedtools |  | RRID:SCR_006646 |  |
| Software, algorithm | STAR v2.5.3a | https://github.com/alexdobin/STAR | RRID:SCR_015899 |  |
| Software, algorithm | Sigmaplot, v12.5 | Systat Software | RRID:SCR_003210 |  |
| Software, algorithm | Custom code for bioinformatics analysis | https://github.com/eric-d-larson/OE_TRPM5 |  |  |
| Software, algorithm | OPM | https://github.com/qi2lab/opm |  |  |

### Overview of the method for transcriptional profiling of low abundance cell populations

For transcriptional profiling of TRPM5-bearing MVC_eGFP cells and OSN_eGFP+ cells that constitute a small fraction of the cells in the epithelium, we used FACS to separate the cell populations targeted for RNAseq (Amamoto et al., 2019). In our experiments, we isolated the cells from mice that expressed fluorescent marker proteins appropriate for cell sorting. OSNs were expressing mCherry under the control of OMP promoter. eGFP was expressed in MVCs and a subset of OSNs (OSN_eGFP+ cells) under control of the TRPM5 promoter.

### Generation of OMP-H2B::Cherry mice

A PacI cassette containing PacI-H2B::mCherry-pA PGK-puro-pA-PacI was inserted into an OMP gene-targeting vector (pPM9)(Mombaerts et al., 1996), which replaces the OMP coding sequence with the PacI cassette and expresses a H2B::mCherry fusion protein. Animals are maintained in a mixed 129/B6 background.

### Animals

Mice with TRPM5-driven eGFP expression (Clapp et al.,2006) were crossed with OMP-H2B::Cherry mice. The TRPM5-eGFP mice and TRPM5 knockout mice(Damak et al., 2006) were obtained with written informed consent from Dr. Robert Margolskee. Lines were maintained separately as homozygous and backcrossed regularly. Experiments were performed on mice from the F1 generation cross of TRPM5-eGFP and OMP-H2B::Cherry mice (OMP-H2B::mCherry/TRPM5-eGFP). PCR was used to verify genotype of experimental mice for eGFP and mCherry expression. Both male and female mice were used for experiments with ages ranging from 3- 8 months. Estrous and cage mate information was collected for all female mice in conjunction with experimental use. Mice were housed in passive air exchange caging under a 12:12 light/dark cycle and were given food and water *ad libitum*. Mice were housed in the National Institutes of Health approved Center for Comparative Medicine at the University of Colorado Anschutz Medical Campus. All procedures were performed in compliance with University of Colorado Anschutz Medical Campus Institutional Animal Care and Use Committee (IACUC) that reviews the ethics of animal use.

In our vivarium we have ventilated cages (HV cages) where air is mechanically exchanged with fresh air once every minute and static cages (LV cages) where air is exchanged passively through a filter in the cover. When we moved the OMP-H2B::mCherry/TRPM5-eGFP to HV cages we noticed a decrease in the number of OSN_eGFP+ cells sorted per mouse (Figure 1-figure supplement 1a,b and c), suggesting that changes in ventilation conditions affect TRPM5 promoter-driven expression of eGFP. Following this observation, mice were moved back to LV cages. We proceeded to study the dependence of the number of OSN_eGFP+ cells sorted on the number of days in LV vs. HV cages. The number of OSN_eGFP+ cells is positively correlated with the number of days the animal spends in LV cages (Figure 1 – figure supplement 1d) and negatively correlated to the number of days the animals spend in the HV cages (Figure 1 – figure supplement 1e). Generalized linear model (GLM) analysis found significant differences for the number of OSN_eGFP+ cells sorted as a function of the number of days in LV cages (p<0.05, 26 observations, 24 d.f., F-statistic = 5.64, p-value for GLM <0.05) and the number of days in HV cages (p<0.05, 26 observations, 24 d.f., F-statistic = 5.99, p-value for GLM <0.05). For RNAseq experiments one FACS sort was done using cells from mice born and maintained in HV housing, and the OSN_eGFP+ yield was low. Subsequently, we performed all FACS with cells isolated from the olfactory epithelium of mice raised in LV cages.

### Tissue dissociation of the olfactory epithelium

Following euthanasia via CO2 inhalation, the olfactory epithelium was immediately removed from the nasal cavity and epithelial tissue was separated from the bone in the turbinates. Care was taken not to include respiratory epithelium. The epithelium was dissociated enzymatically with Dispase II (2 mg/ml) diluted in Ringer’s solution (145mM NaCl, 5mM KCL, 20mM HEPES, 1mM MgCL_2_, 1mM CaCl_2_, 1mM Ny-Pyruvate, 5mM Glucose) (∼25 minutes at 37^0^C) followed by an incubation in a papain plus Ca/Mg++ free Ringer’s solution (Ca/Mg++ free Ringer’s: 145mM NaCl, 5mM KCL, 20mM HEPES, 1mM Ny-Pyruvate, 1mM EDTA, L-cysteine: 1mg L-cysteine /1.5mL Ca/Mg++ free Ringer’s, Papain:1-3ul/1mL Ca/Mg++ free Ringer’s), for ∼40-45 minutes at 37^0^C. Following incubation, DNase I (Promega) at 0.05U/μl and RNAse free 10x Reaction buffer (1:20) were added to solution and the tissue was gently triturated using a ∼1mm opening pipette. Isolated OSNs were collected from supernatants via centrifugation and resuspended in cell sorting medium of 1x PBS (diluted from commercial 10x PBS, pH 7.4) and BrainPhys Neuronal Medium (Stemcell Technologies). Initially, isolated cells were examined with a confocal microscope to confirm efficacy of dissociation methods, and examine cell types and fluorescence. For RNAseq, cells were strained through a 40 μm cell strainer and kept on ice until sorted via flow cytometry.

### Flow cytometry

Fluorescence activated cell sorting was performed in the University of Colorado Cancer Center Flow Cytometry Core on a Beckman Coulter MoFlo Astrios EQ using MoFlo Astrios Summit Software (6.3.1.16945). eGFP signal was detected using a 488 nm laser and a bandpass 526/52nm collection filter. mCherry signal was detected using a 561 nm laser and a bandpass 614/20 nm collection filter. The 488nm laser was also used to detect light scatter. The threshold was set at 3%. Gating was set to exclude doublets and optimized as cell populations emerged based on fluorescent markers. Flow cytometry calibration beads for AcGFP1/eGFP and mCherry (Takara, 632594, 632595) were used as fluorescence intensity controls. Olfactory epithelium cell suspensions from wild type and OMP-H2B::Cherry mice or TRPM5-eGFP mice were sorted as controls for auto fluorescence for eGFP and mCherry populations respectively. Cells were sorted into RNAprotect Tissue Reagent (Qiagen).

### RNA-extraction

Total RNA was extracted from sorted, pooled cells from each cell population using the RNeasy Plus Micro Kit (Qiagen) according to the manufacturers recommended protocol.

### RT-qPCR

Quantitative reverse transcription polymerase chain reaction (RT-qPCR) was used to assess and confirm identities of cell types from each of the sorted cell populations. Following total RNA extraction, RT-qPCR was performed in the PCR core at University of Colorado Anschutz Medical Campus for the following markers: OMP, TRPM5, eGFP and ChAT. Primers and probes used for eGFP, TRPM5 and OMP were described in (Oshimoto et al., 2013). Predesigned primers and probes for ChAT were purchased from Life Technologies. The mRNA for these targets was measured by RT-qPCR using ABI QuantStudio 7 flex Sequence detector. 1μg total RNA was used to synthesize cDNA using the High Capacity c-DNA Reverse Transcription kit (ABI-P/N 4368814). cDNA was diluted 1: 2 before PCR amplification.

The TaqMan probes were 5’labeled with 6-carboxyfluorescein (FAM). Real time PCR reactions were carried out in MicroAmp optical tubes (PE ABI) in a 25 μl mix containing 8 % glycerol, 1X TaqMan buffer A (500 mM KCl, 100 mM Tris-HCl, 0.1 M EDTA, 600 nM passive reference dye ROX, pH 8.3 at room temperature), 300 μM each of dATP, dGTP, dCTP and 600 μM dUTP, 5.5 mM MgCl2, 1X primer-probe mix, 1.25 U AmpliTaq Gold DNA and 5 μl template cDNA. Thermal cycling conditions were as follows: Initiation was performed at 50°C for 2 min followed by activation of TaqGold at 95°C for 10 min. Subsequently 40 cycles of amplification were performed at 95°C for 15 secs and 60°C for 1 min. Experiments were performed with duplicates for each data point. Each PCR run included the standard curve (10 fold serially diluted pooled cDNA from control and experimental samples), test samples, no-template and NORT controls. The standard curve was then used to calculate the relative amounts of targets in test samples. Quantities of targets in test samples were normalized to the corresponding 18s rRNA (PE ABI, P/N 4308310).

### RNA sequencing and pre-processing

RNA quality control, library preparation, and sequencing were performed at the University of Colorado Genomics and Microarray core. Extracted RNA was used as the input for the Nugen Universal Plus mRNA-seq kit (Redwood City, CA) to build stranded sequencing libraries. Indexed libraries were sequenced using an Illumina NovaSEQ6000. Library preparation and sequencing was performed in two batches, separated by gender. 11 female samples were sequenced with an average depth of 37.3 million +/- SD of 6.5 million read pairs, and 25 male samples were sequenced with an average depth of 34.8 million +/- SD of 3.5 million read pairs. Metadata for the samples submitted are shown in Figure 2 - figure supplement 3. Raw BCL files were demultiplexed and converted to FASTQ format. Trimming, filtering, and adapter contamination removal was performed using BBDuk (Bushnell).

### RNA Sequencing Analysis

Transcript abundance was quantified from trimmed and filtered FASTQ files using Salmon v1.2.1(Patro et al., 2017) and a customized Ensembl GRCm38 (release 99) transcriptome (Zerbino et al., 2018). A customized version of the transcriptome was prepared by appending FASTA sequences of eGFP and mCherry to the GRCm38 FASTA file. The corresponding gene transfer format (GTF) file was modified accordingly to incorporate the new transcripts. Transcript abundance was summarized at the gene level using the TxImport (Soneson et al., 2015) package in R. Differential gene expression was quantified using DESeq2 (Love et al., 2014) with default parameters after removing genes with an average count of < 5 reads in each group. Significance was determined by FDR-adjusted p-value < 0.05. TopGO was used for gene ontology analysis (Alexa and Rahnenfuhrer, 2020). The input to TopGO was a list of significant DEGs and a list of all detected genes in the dataset. Enrichment was calculated by dividing the number of detected genes by the number of expected genes within each ontology of the TopGO output. To make the bar graphs in Figures 4 and 5, enrichment scores of downregulated GO terms were multiplied by -1 for visualization. Heatmap visualization was performed using *pHeatmap* in R (Kolde, 2019).

### RNA-sequence data comparison with Ualiyeva et al 2020

Raw counts for this study and for Ualiyeva et al (GEO GSE139014)(Ualiyeva et al., 2020) were converted to log10(reads per million (RPM) +1). These RPM values were used to generate heatmaps to show the expression values of specific transcripts. No quantitative assessment was performed between the two studies.

### Tissue Preparation for Fluorescence Microscopy and *in situ*

For euthanasia, mice were anesthetized with ketamine/xylazine (20–100 _g/g of body weight), perfused transcardially with 0.1 M phosphate buffer (PBS) followed by a PBS-buffered fixative (EMS 32% Paraformaldehyde aqueous solution diluted to 4% with 1x PBS). The nose was harvested and postfixed for 12 h before being transferred for cryoprotection into PBS with 20% sucrose overnight. The olfactory epithelium was cryosectioned coronally into 16 μm -thick sections mounted on Superfrost Plus slides (VWR, West Chester, PA) coated with poly-D-lysine.

### *In situ* followed by immunohistochemistry (IHC)

*In situ* hybridization was performed with the hybridization chain reaction method (Choi et al., 2018) using HCR v3.0 Probe Sets, Amplifiers, and Buffers from Molecular Instruments, Inc. Frozen slides were allowed to thaw and dry, baked at 60°C for 1 hour, then immersed in 70% ethanol overnight at 4°C, and allowed to dry again completely. Slides were inverted and placed on a Plexiglas platform inside a humidified chamber; subsequent steps were performed using this setup. Slides were incubated in 10 μg/μl proteinase K for 15 minutes at 37°C, then pre-hybridized with HCR hybridization buffer (30% formamide buffer from Molecular Instruments) for 30 minutes at 37°C. *Trpm5*-*B3* probes and *OMP-B2* probes (0.8 pmol of each probe in 100 μl HCR hybridization buffer per slide) were added, and slides were hybridized overnight at 37°C. Slides were briefly incubated in undiluted HCR Wash Buffer (30% formamide buffer from Molecular Instruments) for 20 minutes at 37°C. Excess probes were removed by incubating slides for 20 minutes each at 37°C in solutions of 75% HCR Wash Buffer / 25% SSCT (5X SSC, 0.1% Tween, diluted in RNAse free water), 50% Buffer / 50% SSCT, 25% Buffer / 75% SSCT, and 100% SSCT. Slides were incubated in 100% SSCT at room temperature for 20 minutes, then in Amplification Buffer (Molecular Instruments) at room temperature for 1 hour. B3 hairpins labeled with Alexa Fluor 647 and B2 hairpins labeled with Alexa Fluor 546 were prepared (12 pmol of each hairpin were heat shocked, then cooled for 30 minutes, and added to 200μl of Amplification Buffer) added to slides, and incubated overnight at room temperature. Excess hairpins were removed with four washes (20 minutes) in SSCT at room temperature. Slides were then processed with IHC protocol to stain for GFP. At room temperature, tissue was permeabilized with Triton X-100 0.1% in PBS for 30 minutes, washed three times with PBS, blocked with Donkey serum 5% and Tween 20 0.3% in PBS for 1 hour, incubated with Chicken anti-GFP primary antibody (1:500 in blocking solution, AB_2307313 Aves labs) overnight, washed three times with PBS and incubated with Donkey anti-Chicken secondary antibody conjugated with alexa fluor 488 (1:500 in blocking solution, 703-545-155 Jackson ImmunoResearch laboratories).After three final washes with PBS, slides were mounted using Fluoromount-G™ mounting medium with DAPI (Thermo Fisher Scientific).

### Confocal fluorescence microscopy

Microscopy was performed with confocal microscopes (Leica SP8, Nikon A1R or 3i Marianas).

### Three-dimensional tissue imaging

For three-dimensional imaging, a high numerical aperture (NA) oblique plane microscope was used (Dunsby, 2008; Sapoznik et al., 2020). Briefly, this variant on a light sheet microscope only uses one objective to interface with the sample. The sample is illuminated from the epi-direction using an obliquely launched light sheet. Emitted fluorescence is detected through the same primary objective used for illumination. A secondary and tertiary objective optically resample the emitted fluorescence to image the fluorescence resulting from the obliquely launched light sheet onto a detector(Dunsby, 2008; Sapoznik et al., 2020). For the primary, secondary, and tertiary objectives we used a high-NA silicone immersion objective (Nikon ×100 NA 1.35, 0.28–0.31□mm working distance), a high-NA air immersion objective (Nikon ×40 NA 0.95, 0.25–0.16□mm working distance), and a bespoke glass-tipped objective (AMS-AGY v1.0, NA 1.0, 0□ mm working distance), respectively. Images were acquired by a high-speed scientific CMOS camera (Photometrics Prime BSI) using custom Python software ((Sapoznik et al., 2020), https://github.com/qi2lab/opm).

The obliquely launched light sheet was set to 30 degrees above the coverslip. The sample was translated in one lateral dimension (x) at a constant speed by a scan optimized stage. The scan speed was set so that images with a 5-millisecond exposure time were acquired at 200 nm spacing over a distance of 5.5 mm. This constant speed scan was performed for the same volume, cycling through three excitation wavelengths (405, 488, 635 nm) and three sample height positions (z), with 20% overlap. Once the cycle of wavelengths and height positions completed, the sample was then laterally displaced (y), again with a 20% overlap, and the scan was repeated over a 5.5 mm x 5.5 mm x .035 mm (x,y,z) imaging volume. Raw data was orthogonally deskewed, stitched, and fused using custom Python code and BigStitcher(Hörl et al., 2019). After export, each inset image was deconvolved using Microvolution and measured point spread functions.

### Statistical analysis

Statistical analysis was performed in Matlab (Mathworks, USA). Statistical significance was estimated using a generalized linear model (GLM), with post-hoc tests for all data pairs corrected for multiple comparisons using false discovery rate (Curran-Everett, 2000). The post hoc comparisons between pairs of data were performed either with a t-test, or a ranksum test, depending on the result of an Anderson-Darling test of normality. 95% CIs shown in the figures as vertical black lines or shading bounding the lines were estimated by bootstrap analysis of the mean by sampling with replacement 1000 times using the bootci function in MATLAB.

**Table 1.** Levels of expression and adjusted p-value for the olfactory receptor genes whose levels are significantly higher in OSN_eGFP+ compared to OSN_eGFP-. These olfactory receptors had an adjusted p-value for expression level difference between OSN_eGFP+ compared to OSN_eGFP- and had a fold change > 4 and average expression > 100 counts.

| Name | OSN_eGFP- | OSN_eGFP+ | MVC_eGFP | p-value adjusted |
| --- | --- | --- | --- | --- |
| *Olfr292* | 3.61 | 959 | 4.29 | 6.35E-09 |
| *Olfr282* | 2.05 | 486 | 0 | 8.01E-05 |
| *Olfr1434* | 53.3 | 7730 | 0 | 9.71E-16 |
| *Olfr390* | 101 | 10800 | 43.1 | 1.56E-16 |
| *Olfr305* | 6.14 | 612 | 0 | 6.3E-12 |
| *Olfr293* | 6.96 | 664 | 16.9 | 1.42E-07 |
| *Olfr378* | 3.41 | 322 | 0 | 1.1E-06 |
| *Olfr128* | 39.6 | 3660 | 12.7 | 7.33E-14 |
| *Olfr344* | 16.2 | 1050 | 0 | 1.4E-11 |
| *Olfr307* | 7.59 | 393 | 0 | 3.76E-06 |
| *Olfr391* | 156 | 8000 | 9.79 | 1.01E-15 |
| *Olfr299* | 13.1 | 651 | 0 | 4.58E-09 |
| *Olfr142* | 36.4 | 1720 | 3.77 | 1.62E-08 |
| *Olfr1* | 147 | 5720 | 52.7 | 3.08E-10 |
| *Olfr1279* | 16.4 | 552 | 10.9 | 3.52E-07 |
| *Olfr39* | 13.8 | 388 | 21 | 2.81E-06 |
| *Olfr1447* | 64.1 | 1610 | 0 | 1.23E-07 |
| *Olfr728* | 2150 | 45700 | 320 | 6.13E-22 |
| *Olfr727* | 560 | 11000 | 179 | 1.64E-07 |
| *Olfr1555-ps1* | 10.1 | 175 | 0 | 0.0397 |
| *Olfr346* | 27.6 | 465 | 0 | 3.85E-05 |
| *Olfr1228* | 533 | 5320 | 35.4 | 3.09E-08 |
| *Olfr1181* | 87.4 | 766 | 4.61 | 0.000766 |
| *Olfr943* | 97.1 | 844 | 2.89 | 0.000886 |
| *Olfr298* | 60.1 | 509 | 0 | 0.00132 |

## Declarations

### Ethics approvals

Mouse experiments were carried out under guidelines of the National Institutes of Health in compliance with University of Colorado Anschutz Medical Campus Institutional Animal Care and Use Committee (IACUC).

### Consent for publication

Not applicable.

### Availability of data and materials

All data sequencing data are available in NCBI SRA https://www.ncbi.nlm.nih.gov/sra/PRJNA632936. The code used for bioinformatics analysis is found in https://github.com/eric-d-larson/OE_TRPM5

### Authors’ contributions

D.R., B.D.B., M.N. and V.R. conceptualized the project. B.D.B. performed FACS, qPCR and RNAseq experiments. E.D.L. performed genomic analysis. P.F. generated the OMP-H2B::Cherry mice. D.S. designed and analyzed in situ experiments. M.N., A.N.B. and C.N. designed experiments. C.N. and J.H.Jr. performed experiments. M.L. performed in situ experiments and literature search and wrote the section on viral infection in the discussion. All authors contributed to writing and editing the manuscript.

## Acknowledgements

We would like to acknowledge the support of Nicole Arevalo for laboratory support, Jerome Artus and Anna-Katerina Hadjantonakis for the construction of the targeting vector and production of OMP-H2B::Cherry mice, Emily Liman for providing TRPM5 and TRPC2 antibody and Catherine Dulac for providing tissue from TRPC2 knockouts.

## Supplemental Information

**Figure 1. figure supplement 1.**
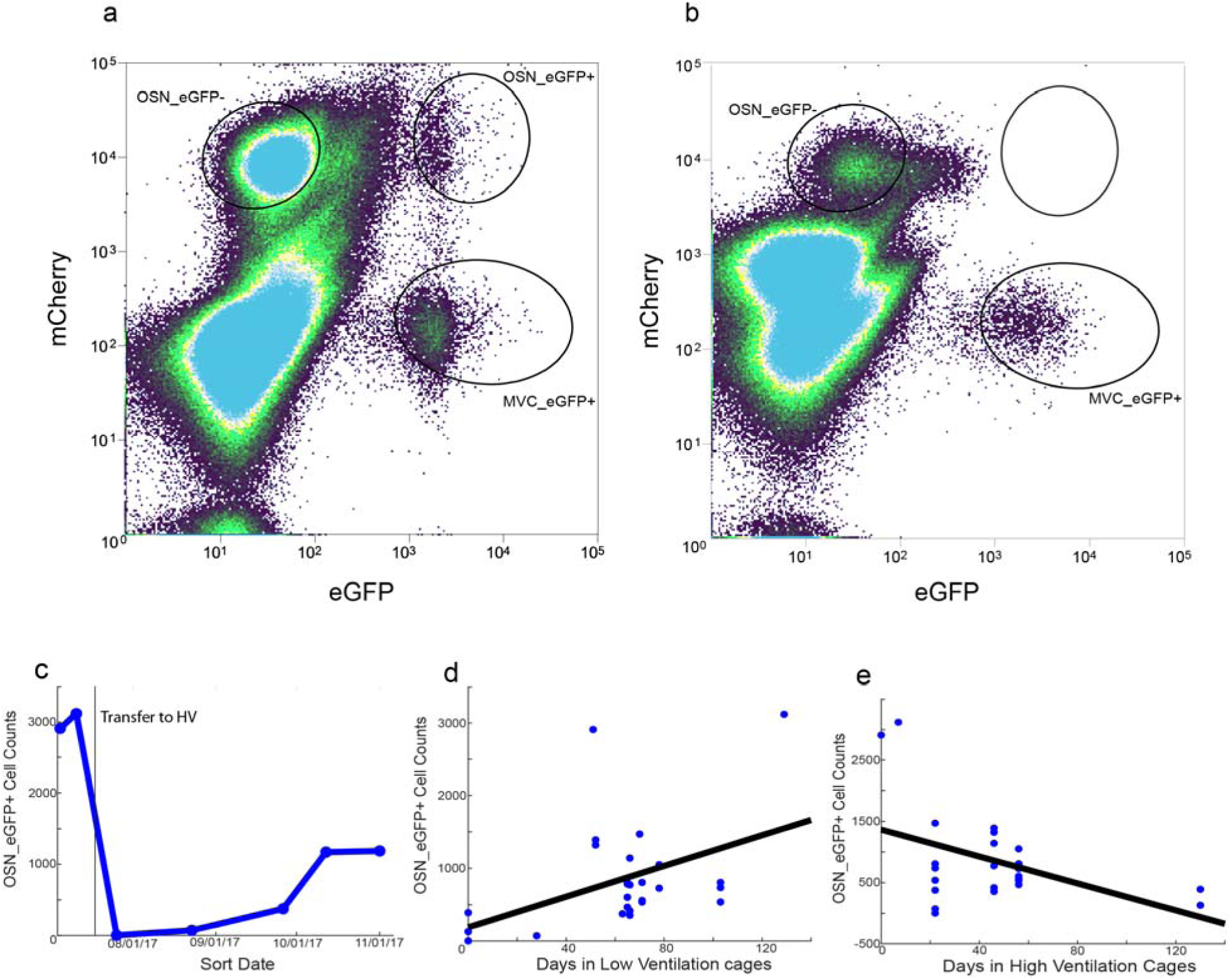
Decreased yield of OSN_eGFP+ cells when mice are moved from low ventilation (LV) to high ventilation (HV) cages. **a and b.** Distribution of mCherry and eGFP fluorescence intensity for FACS-sorted cells that either were not transferred to HV cages (**a**) or were transferred to HV cages for 22 days before sorting (**b**). **c.** Time course showing change in the number of sorted OSN_eGFP+s after mice were transferred to HV cages. **d and e.** Dependence of the yield of OSN_eGFP+ cells after sorting on the number of days in LV cages (**d**) or the number of days in HV cages (**e**).

Figure 2 – figure supplement 1. Excel worksheet with the results of comparison of gene transcription between MVC_eGFP and OSN_EGFP-.

Figure 2 - figure supplement 2. Significant differences in gene ontology for MVC_eGFP+ compared to OSN_eGFP-.

Figure 2 - figure supplement 3. Metadata for the RNAseq.

**Figure 2 – figure supplement 4.**
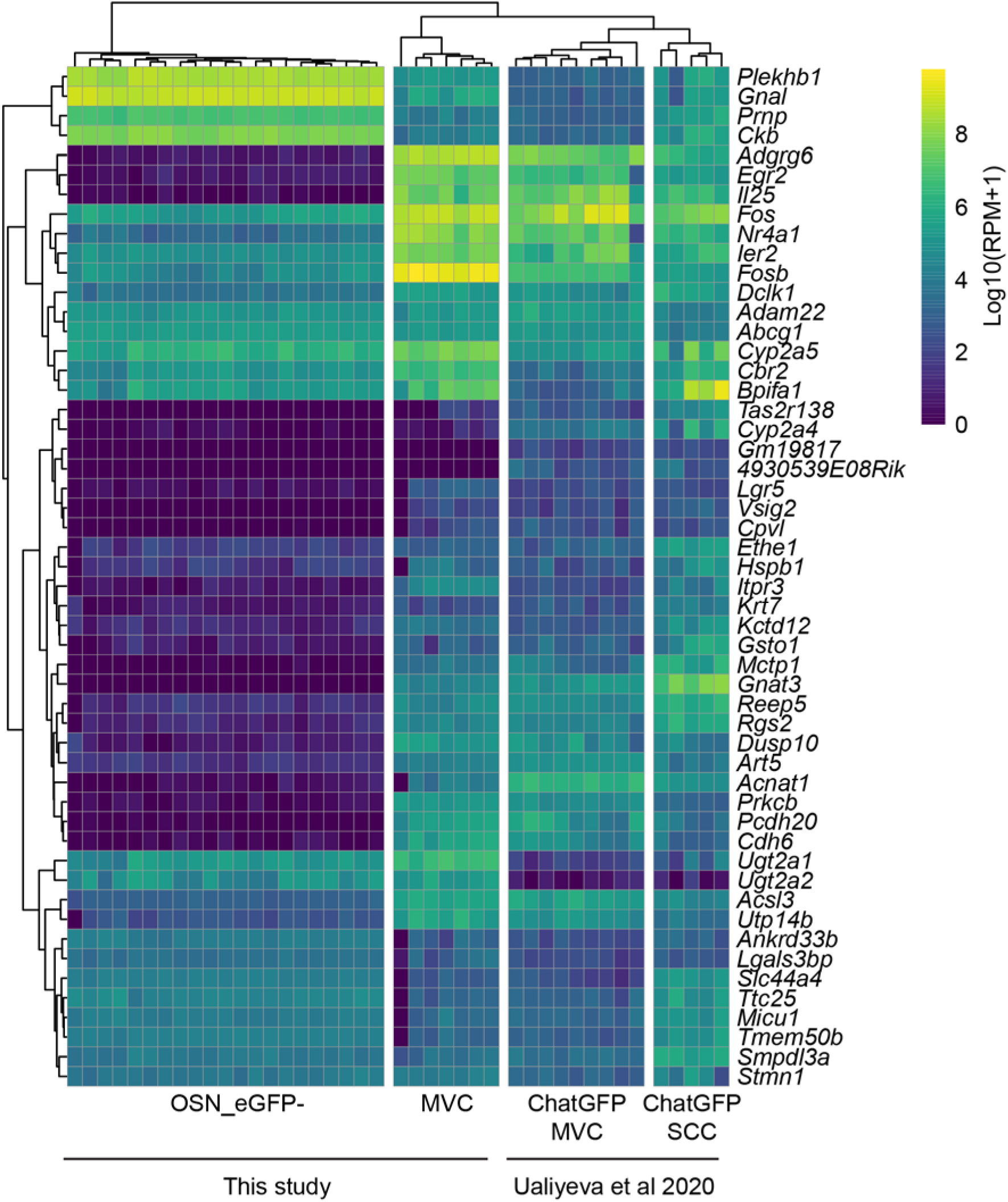
Comparisons of gene expression between MVC_eGFP and OSN_eGFP- cells of this study and ChAT-eGFP MVCs and ChAT-eGFP SCCs profiled in the respiratory epithelium in the study of Ualiyeva and co-workers (Ualiyeva et al., 2020). This comparison is of limited value due to the fact that the gene profiling was performed in two separate studies.

**Figure 3- figure supplement 1.**
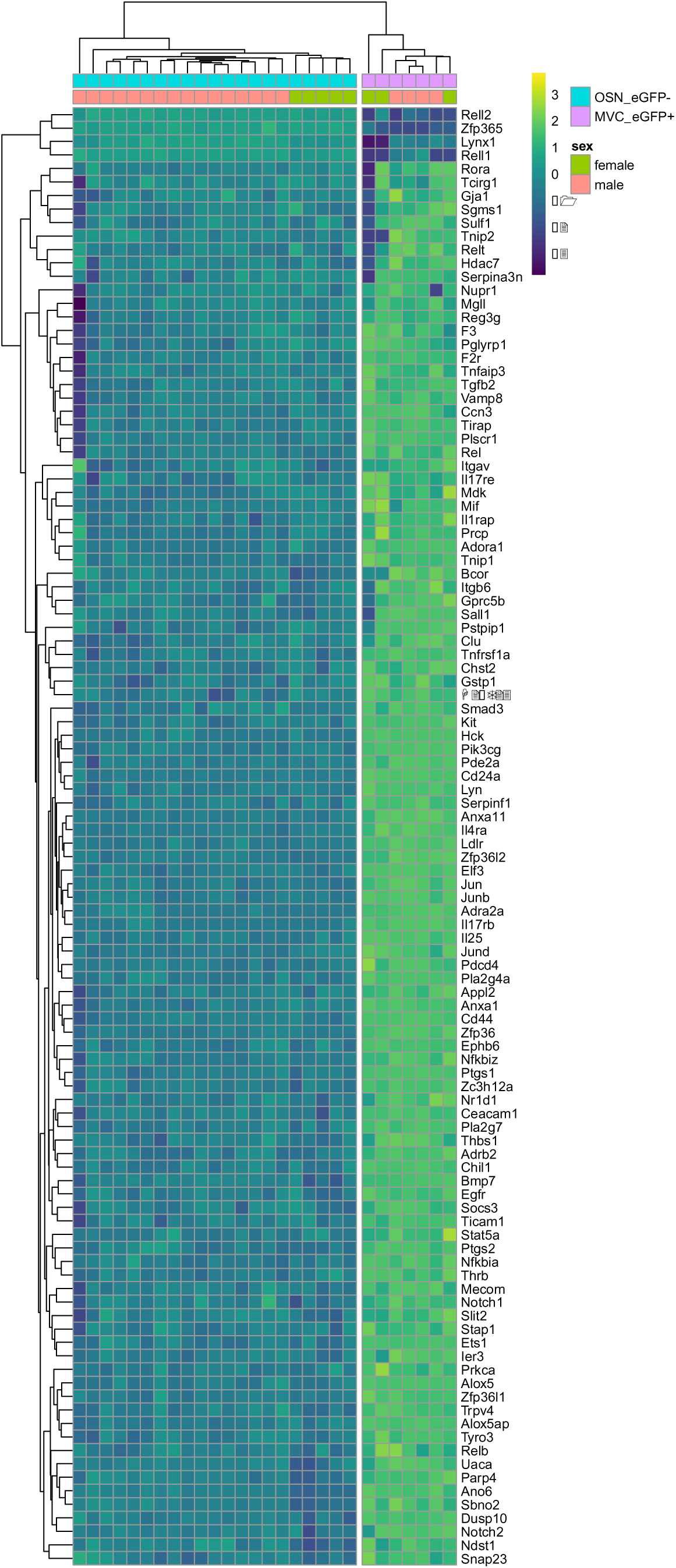
Significant differences in inflammation gene ontology for MVC_eGFP+ compared to OSN_eGFP-.

**Figure 3- figure supplement 2.**
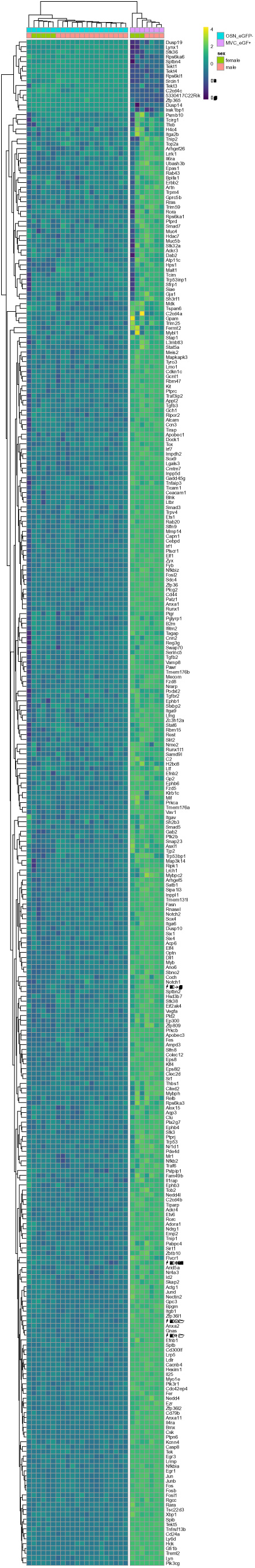
Significant differences in immmunity gene ontology for MVC_eGFP+ compared to OSN_eGFP-.

Figure 4 – figure supplement 1. Excel worksheet with the results of comparison of gene transcription between OSN_EGFP+ and OSN_EGFP-.

**Figure 4 – figure supplement 2.**
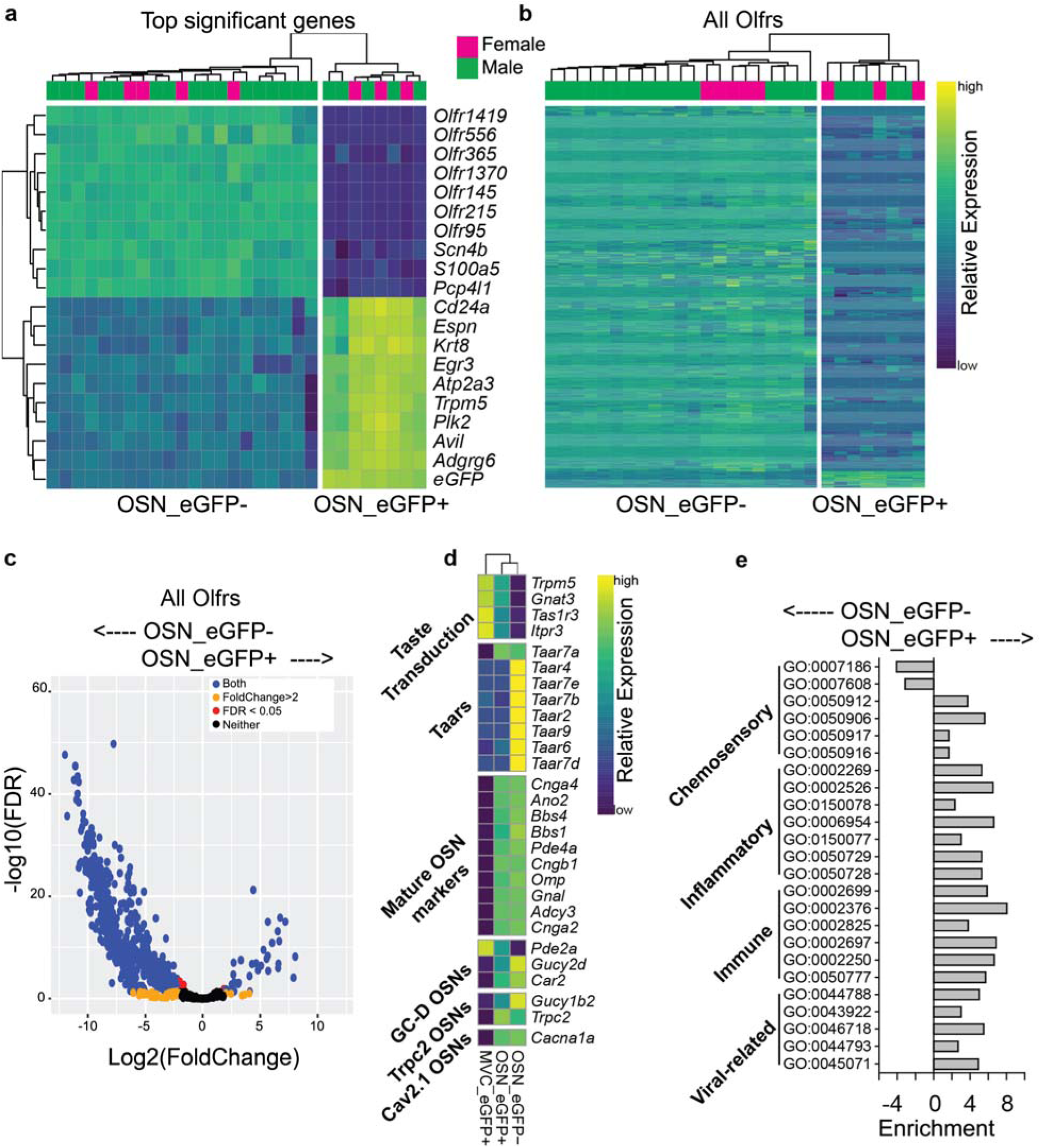
RNAseq comparison of OSN_eGFP- and OSN_eGFP+ cells. **a.**Heatmap showing the top 10 upregulated and top 10 downregulated genes identified by DESeq2. **b.** Heatmap showing all *Olfr* genes detected in the data. For both a and b, row and column order were determined automatically by the *pHeatmap* package in R. For each data point relative expression was calculated by subtracting the average row value from each individual value. **c.** Volcano plot of all Olfactory receptors, demonstrating the small number of enriched olfactory receptors in the OSN_eGFP+ population. **d.** Hierarchical clustering of transcripts for taste transduction and transcripts expressed in canonical and non-canonical OSNs identified by RNAseq as significantly different in expression between the cell groups. We compared expression of transcripts involved in taste transduction, canonical olfactory transduction, and non-canonical OSNs. The non-canonical OSNs considered here included guanilyl-cyclase D (GC-D) OSNs (Juilfs et al., 1997), Trpc2 OSNs (Omura and Mombaerts, 2014), Cav2.1 OSNs (Pyrski et al., 2018), and OSNs expressing trace amine-associated receptors (Taars) (Liberles, 2015). Transcripts identified by DESeq2. **e.** Gene ontology (GO) term enrichment was calculated from differentially expressed genes using *TopGO* in R. An enrichment value for genes with Fischer p value <0.05 was calculated by dividing the number of expressed genes within the GO term by the number expected genes (by random sampling, determined by *TopGO*).

Figure 4 - figure supplement 3. Excel worksheet with the results of comparison of gene transcription between MVC_eGFP cells and OSN_eGFP+.

**Figure 4 figure supplement 4.**
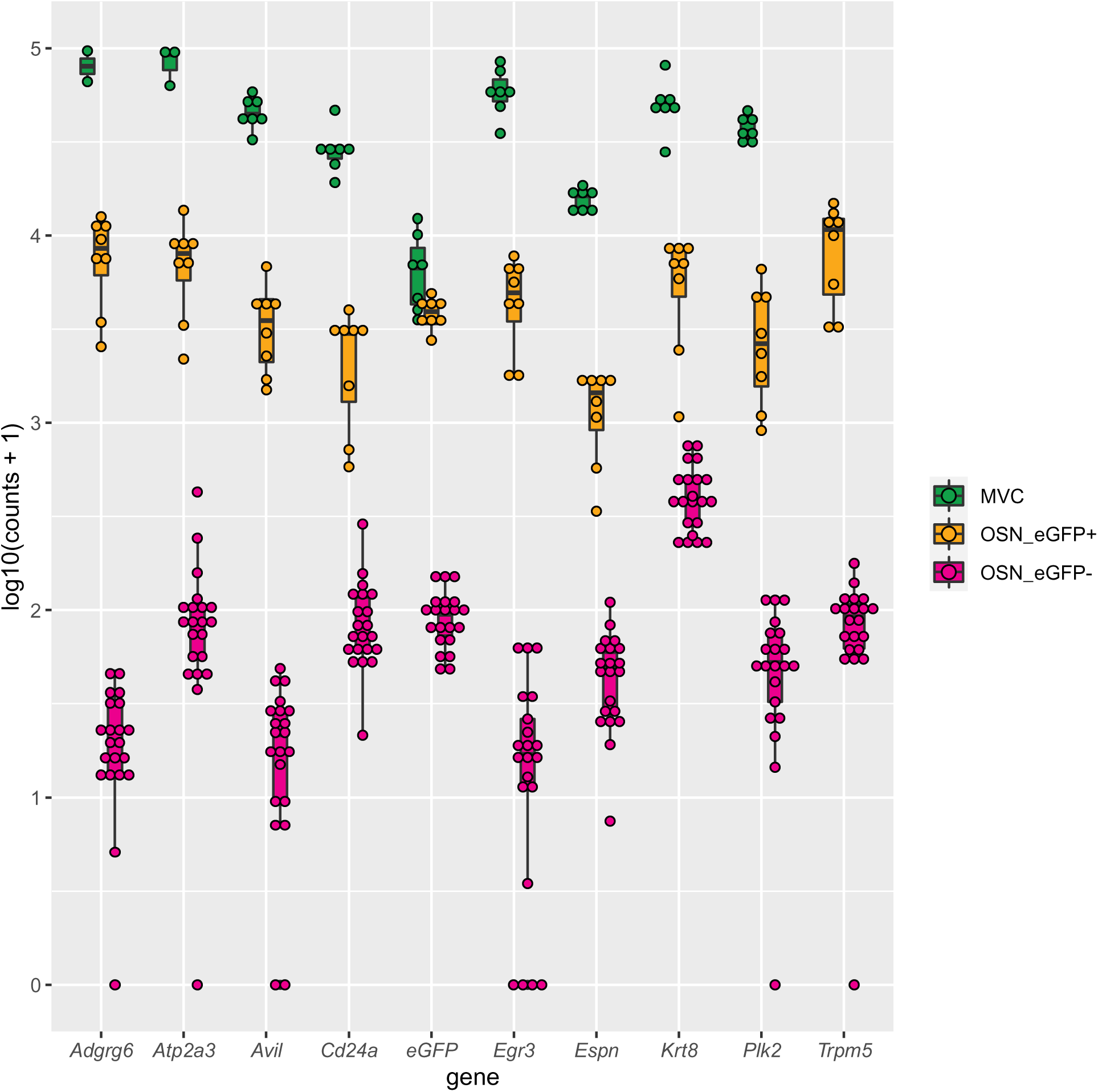
Boxplot showing expression levels for the top 10 genes that are significantly higher in OSN_eGFP+ vs. OSN_eGFP-.

Figure 4 - figure supplement 5. Excel worksheet with the results of genes whose transcription levels were significantly higher in OSN_eGFP+ compared to both MVC_eGFP cells and OSN_eGFP+.

Figure 4 figure supplement 6. GOnet analysis for the 80 genes whose transcription levels were significantly higher in OSN_eGFP+ compared to both MVC_eGFP cells and OSN_eGFP+. https://tools.dice-database.org/GOnet/job0c0357f0-d489-467d-adb7-9d33b52ff850/result

**Figure 4 – figure supplement 7.**
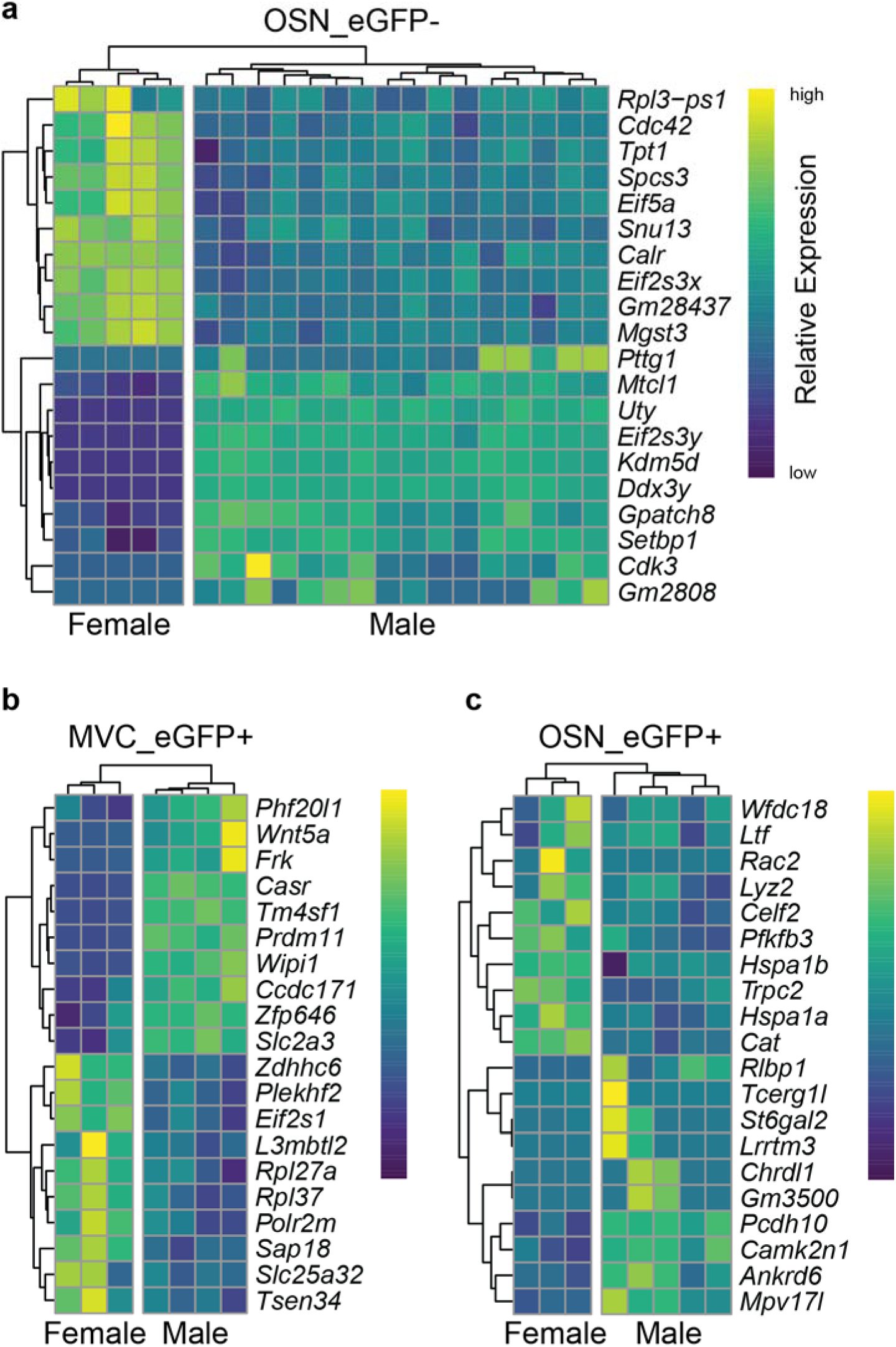
Hierarchical clustering of transcripts identified by RNAseq as significantly differentially expressed between male and female. a. **OSN_eGFP-.** b. MVC_eGFP cells **c.** OSN_eGFP+. Transcripts identified by DESeq2.

**Figure 4 – figure supplement 8.**
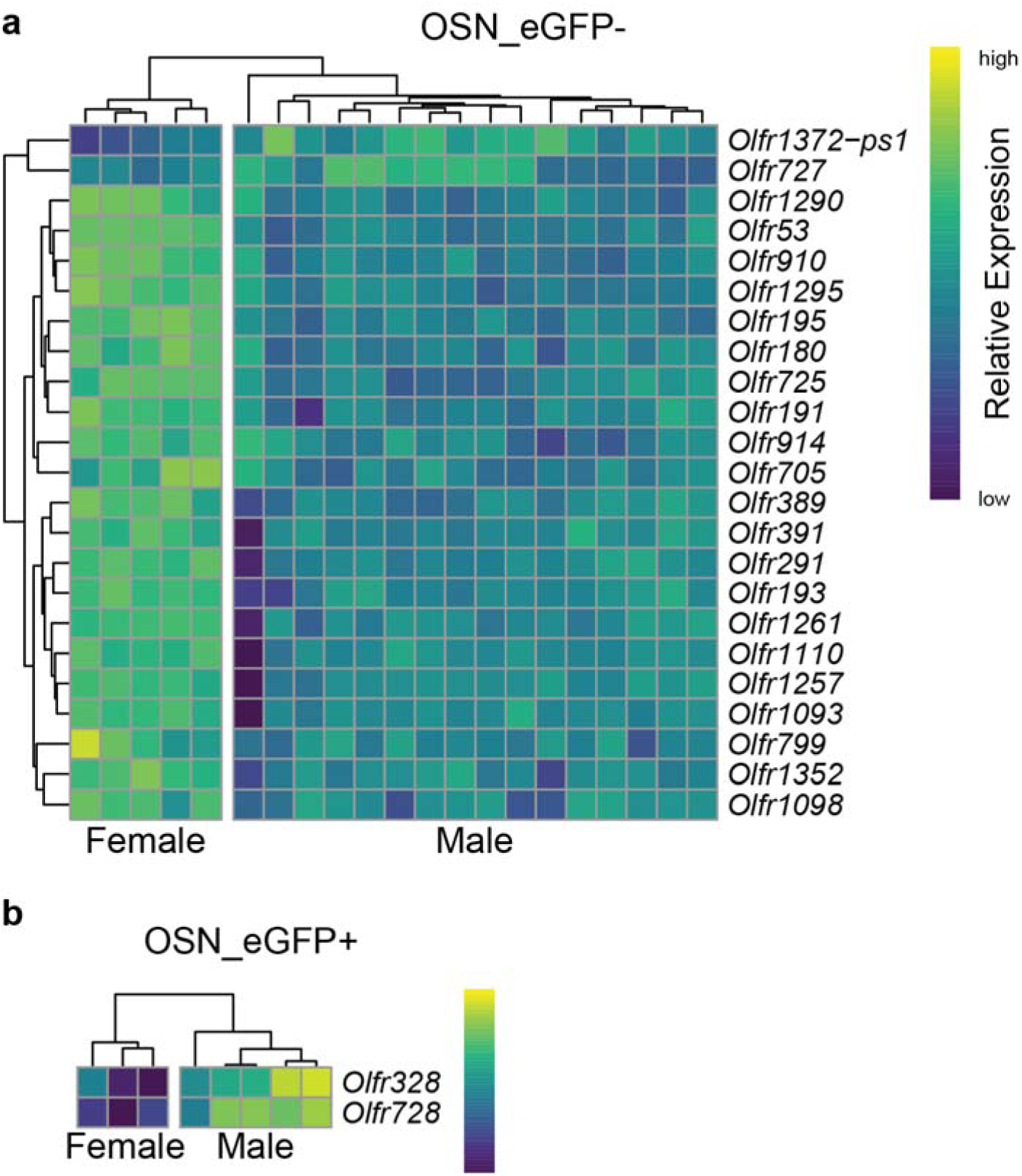
Hierarchical clustering of olfactory receptor transcripts identified by RNAseq as significantly differentially expressed between male and female. **a.** OSN_eGFP-. **b.** OSN_eGFP+. Transcripts identified by DESeq2.

**Figure 5 – figure supplement 1. Movie showing 3D rendering of HCR v3.0 *in situ* hybridization for TRPM5.** Images were acquired with a high numerical aperture (NA) oblique plane microscope (Sapoznik et al., 2020). *In situ* is shown for TRPM5 (yellow) and OMP (green). DAPI is in blue.

