## Supplementary figures and images for "Transcriptional profiling reveals potential involvement of microvillous TRPM5-expressing cells in viral infection of the olfactory epithelium"

### Figure 1, figure supplement 1

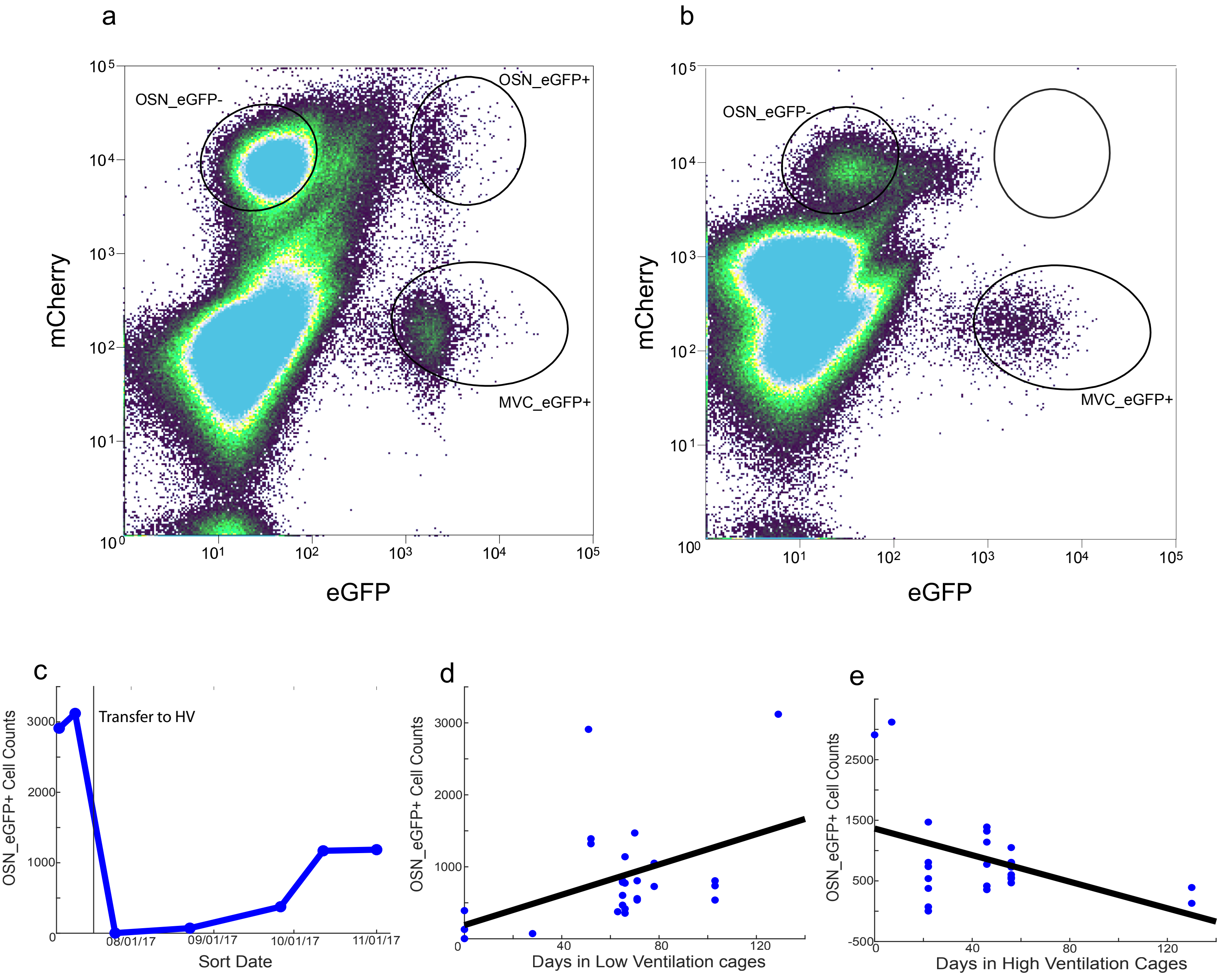

### Figure 3- figure supplement 1

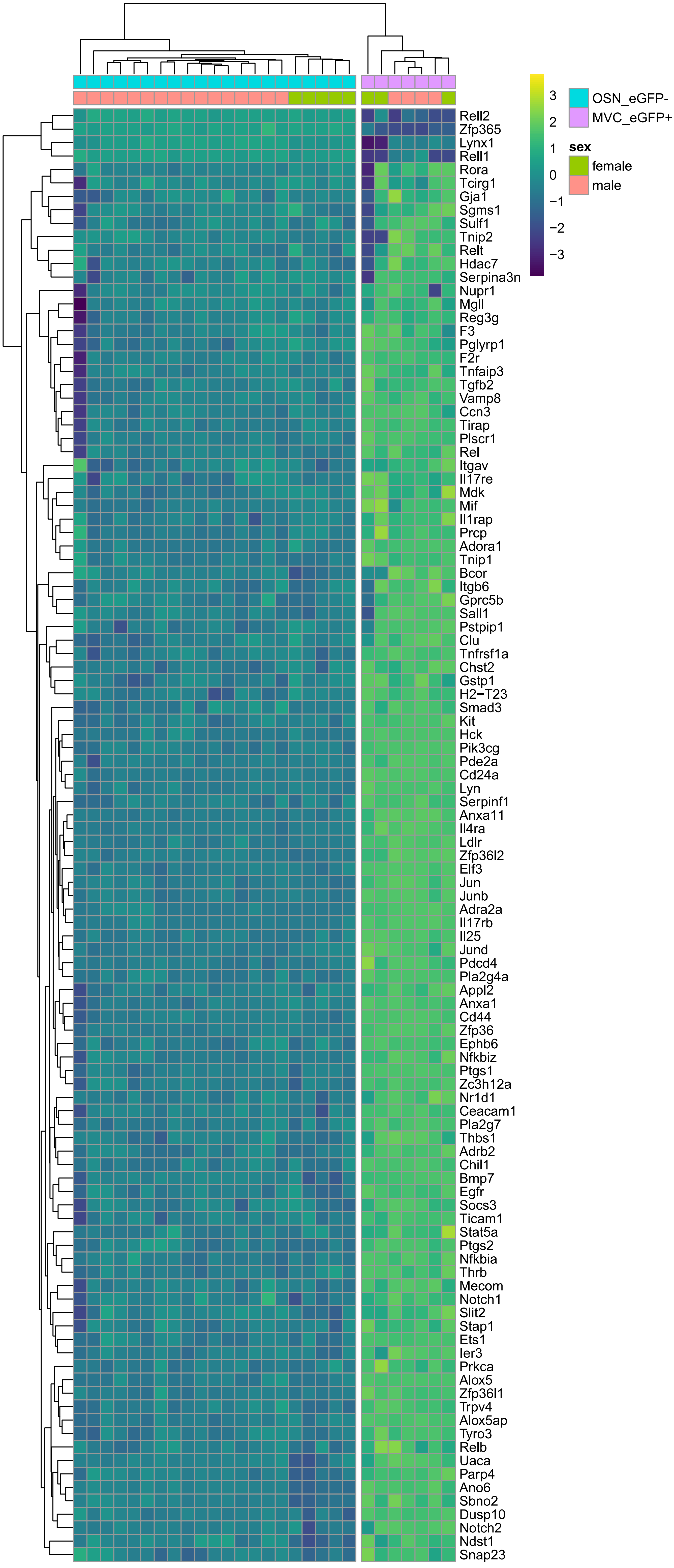

### Figure 3- figure supplement 2

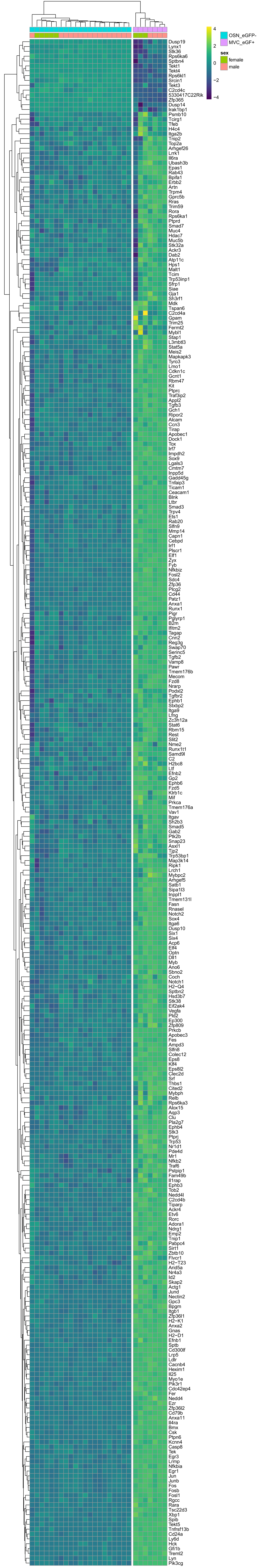

### Figure 4 figure supplement 4

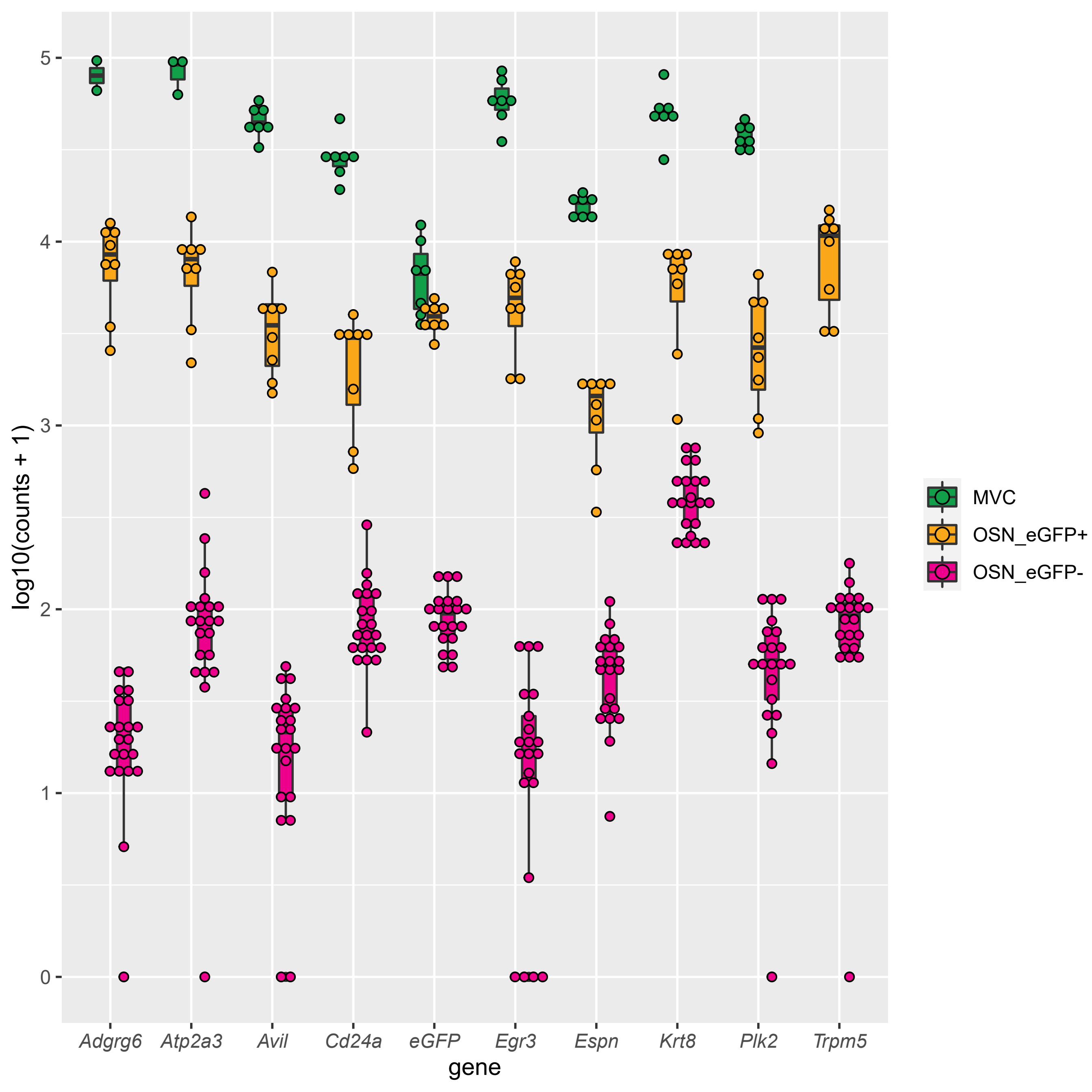

### Supplemental Data 2

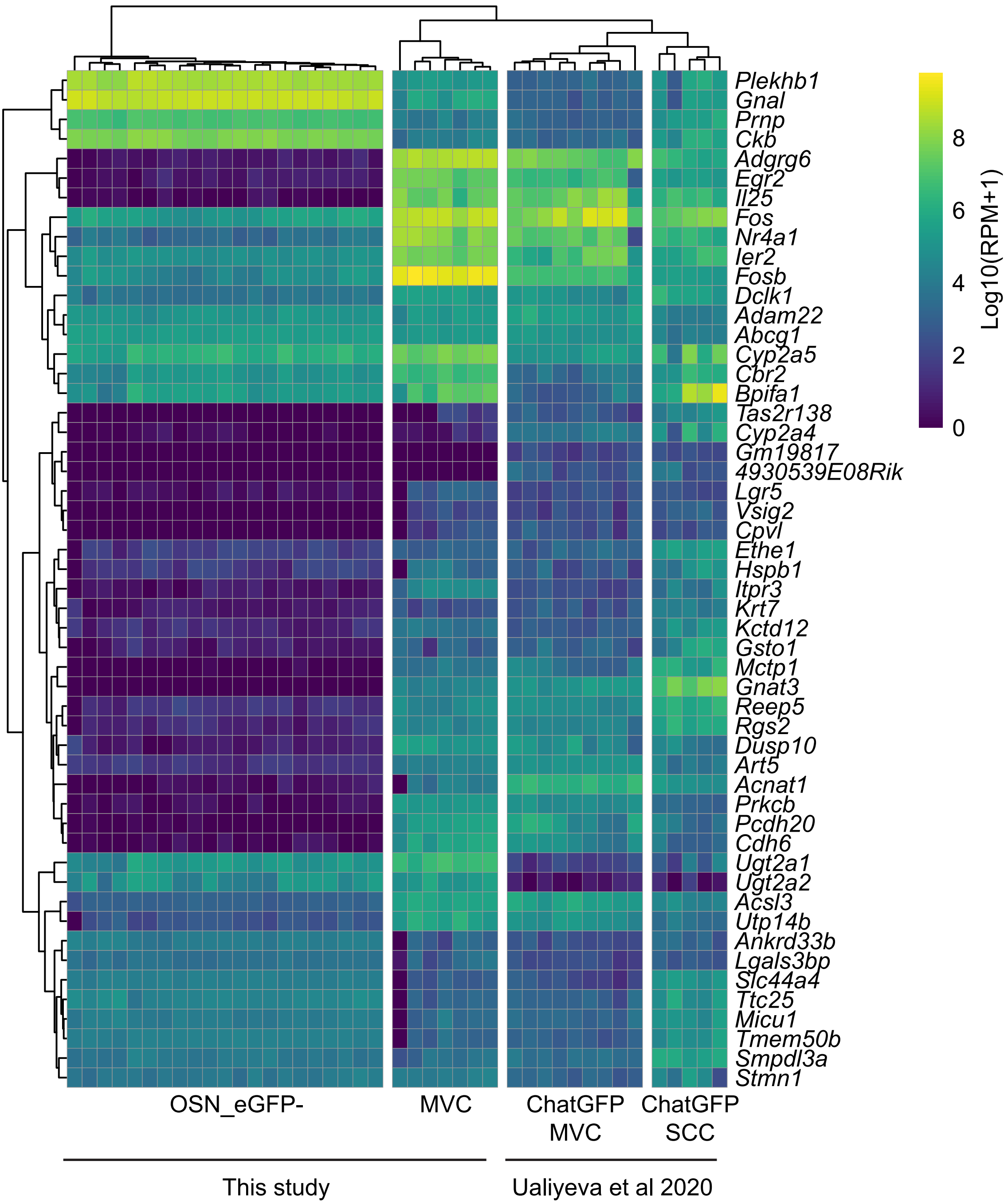

### Supplemental Data 4

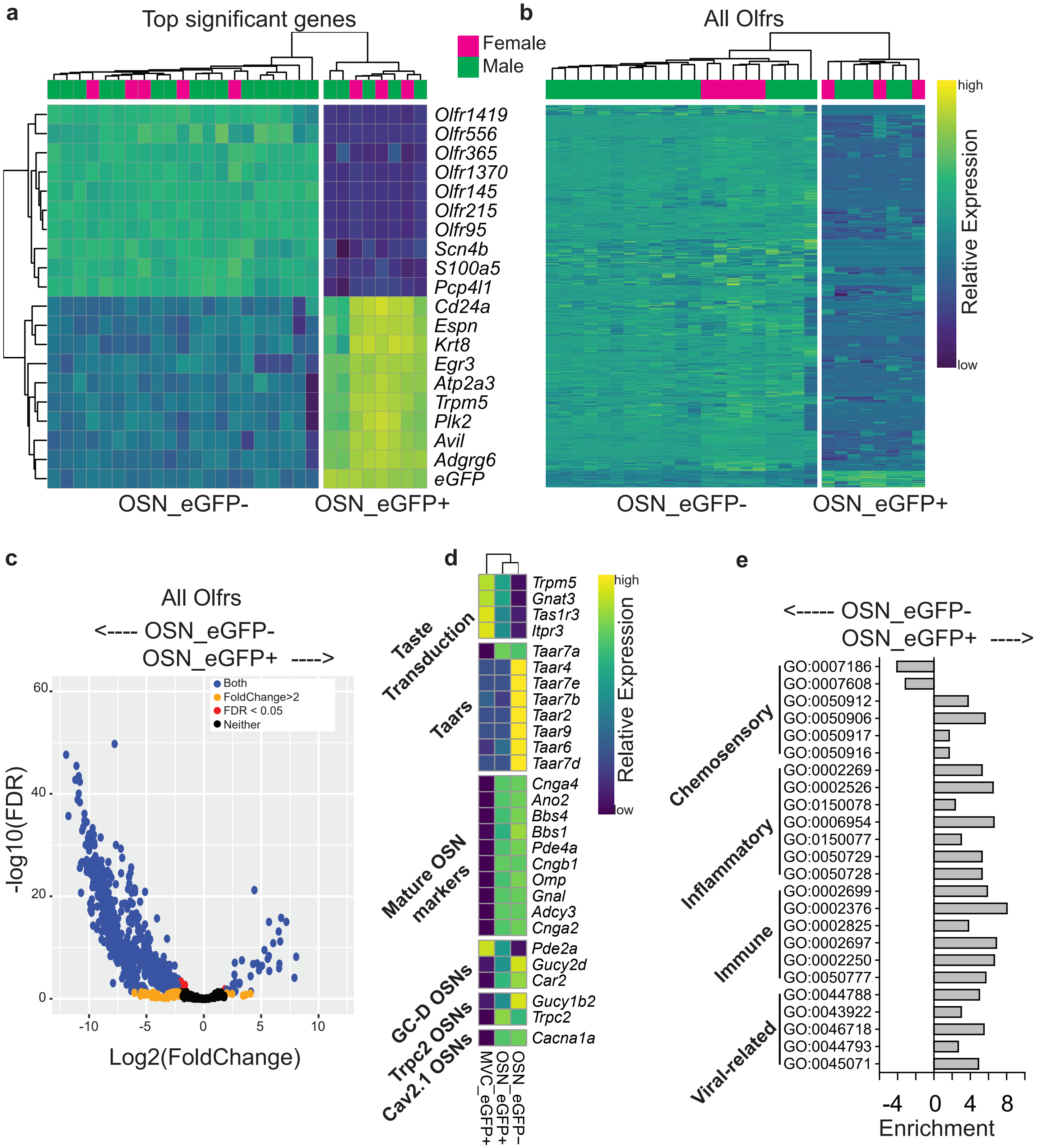

### Supplemental Data 5

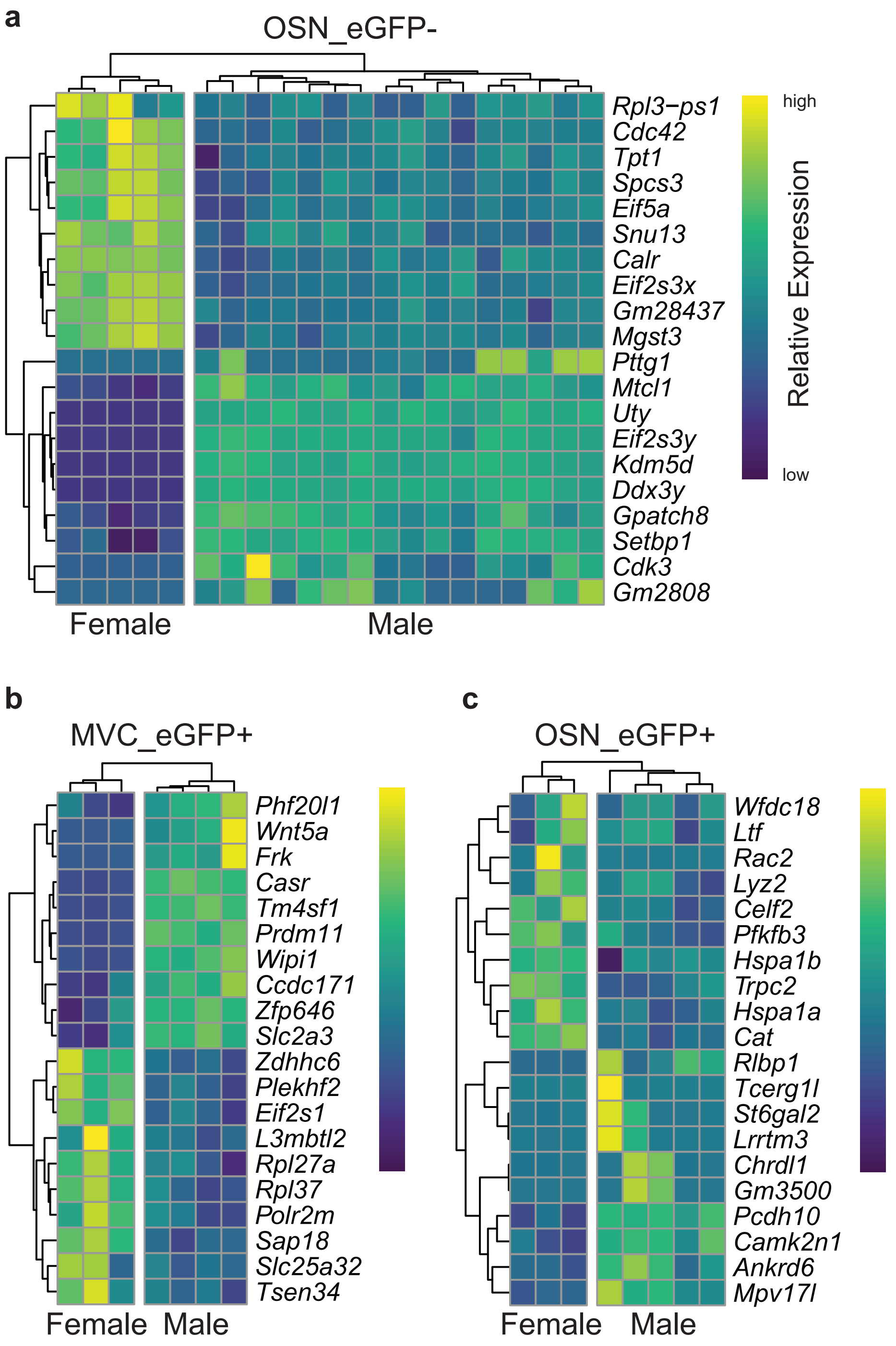

### Supplemental Data 6

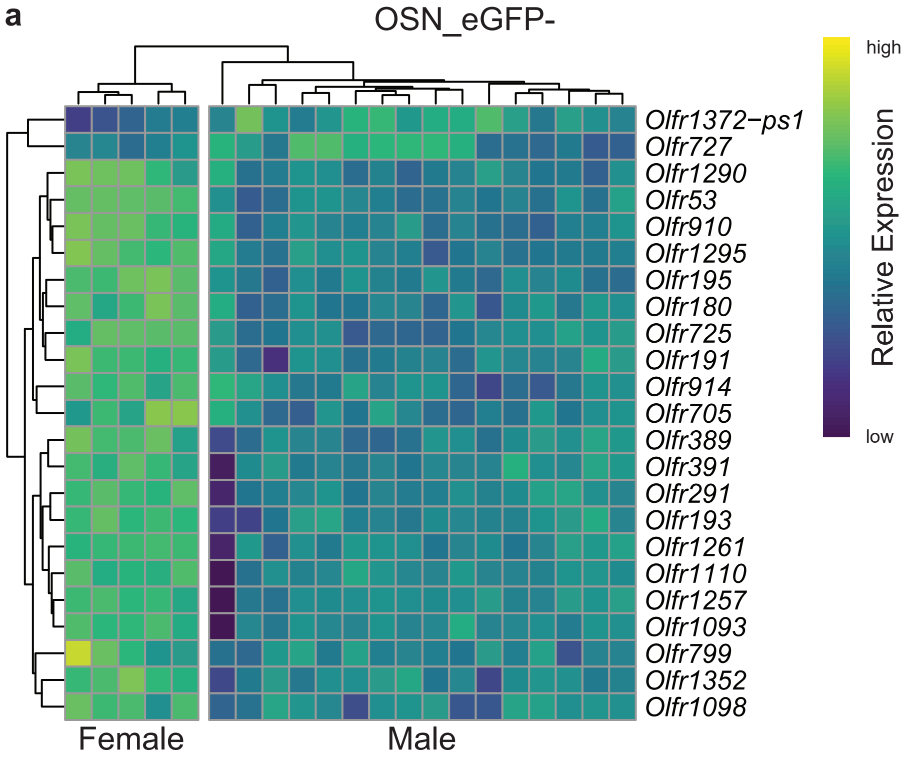

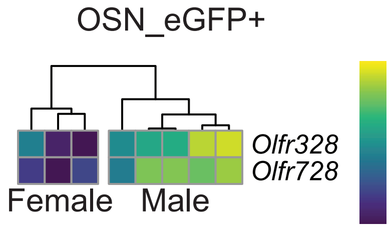
